# Mesenteric ischemia and bacterial translocation precipitate the intoxication phase of yellow fever

**DOI:** 10.1101/2025.03.05.641677

**Authors:** Mateus V. Thomazella, Xueer Qiu, Cássia Gisele Terrassani Silveira, Carolina A. Correia, Mariana P. Marmorato, Erik Sanson, Andrew Norton, Mark Sharobim, Yeh-Li Ho, Marisa Dolhnikoff, Eduard Matkovic, Esper G. Kallas, Amaro Nunes Duarte-Neto, Adam L. Bailey

## Abstract

Yellow fever (YF) is classically conceptualized as a hepatotropic disease; indeed, the liver is the primary site of yellow fever virus (YFV) replication. However, circumstantial evidence suggests that extra-hepatic disease may be important for the ∼30% of YF cases that progress to the severe “intoxication” phase of the disease. Using a Syrian hamster-adapted (HA)-YFV, we worked backwards from observations in humans to examine early events that precipitate the intoxication phase of YF. HA-YFV caused severe disease in ∼80% of infected animals characterized by lethargy and weight loss that progressed to widespread petechiae and death by day 6. Clinical chemistry, coagulation testing, histology, immunohistochemistry, and in-situ hybridization were consistent with a cascade of hepatocyte-specific virus replication causing liver damage and a defect in clotting factor synthesis. Despite a lack of extra-hepatic HA-YFV replication, severe pathology was observed in the intestines and pancreas. Histopathological analysis over the time-course of HA-YFV infection revealed an ischemic pattern in these tissues, culminating in fibrinoid/coagulative necrosis of these organs. Further investigation showed that ischemia-induced erosion of the gut epithelial barrier serves as an entry point for luminal bacteria that spread systemically via the portal system. Thus, the intoxication phase of YF is a sepsis-like syndrome caused by translocation of bacteria from a damaged gastrointestinal tract. Evaluation of human YF cases for these previously overlooked disease features confirmed this overarching mechanism: bacteria were identified in the portal vein and liver parenchyma of fatal YF cases along with elevations in plasma markers of bacteremia and a bacteria-driven inflammatory response. Importantly, blood concentrations of the gastrointestinal damage marker intestinal fatty acid binding protein (I-FABP) were significantly elevated in fatal YF cases relative to non-fatal cases, suggesting that I-FABP measurements could be useful in prognosis and treatment decision making. Our findings tie together several recent and historically unexplained observations surrounding the highly-lethal intoxication phase of YF in humans: a high AST/ALT ratio, “black vomit,” pancreatitis, and paradoxical neutrophilia. A better appreciation for the drivers of mesenteric ischemia, and preemption of bacterial sepsis, may improve outcomes in cases of severe YF.

## INTRODUCTION

Yellow fever virus (YFV) is a positive-sense single-stranded RNA virus with a lipid envelope (Family: *Flaviviridae*; Genus: *Orthoflavivirus*). In addition to being the prototypical orthoflavivirus from which the genus and family “*flavi*” (Latin for “yellow”) name is derived, YFV remains a significant human pathogen (*1–3*). YFV is endemic to equatorial Africa and South America, where “sylvatic cycles” among nonhuman primates and forest-dwelling mosquitoes (e.g., *Haemagogus* and *Sabethes* mosquitoes) maintain a reservoir for zoonotic transmission to humans. Spillover into humans can facilitate a switch to the *Aedes aegypti* mosquito vector that has a preference for feeding on humans, resulting in outbreaks that can threaten larger population centers in an “urban cycle” of transmission. The burden of yellow fever (YF) disease is challenging to measure, but modeling suggests that ∼130,000 severe cases and ∼78,000 deaths occur annually in Africa alone (*4, 5*). South America has seen an expansion of YFV activity beyond its historical river basin regions, with recent outbreaks threatening urban centers (*e.g*., São Paulo 2017-2019) (*6*). Although a highly-effective YF vaccine is available, several factors complicate its implementation, including: long-established egg-based production technology resulting in vaccine shortages; a long list of vaccine contraindications and the risk of severe complications that render many demographics ineligible; and the vast and rugged geography that hinders healthcare delivery in much of YFV’s area of endemicity (*7*). Thus, an estimated 400 million people living in YF-endemic areas are in need of YF vaccination in order to achieve the herd immunity threshold of 80% required to prevent the urban spread of YF (*8*). The shortage of YF vaccines, coupled with the expanding range of *A. aegypti*, is concerning for the emergence (or re-emergence) of sylvatic and urban YF in new and poorly-prepared places (*5*). While this may include occasional season-dependent outbreaks of YF in North America and Europe, spread to Asia or Oceania could lead to the establishment of permanent YFV reservoirs in wild primates, putting large populations of YFV-naïve humans at risk (*8–11*).

The course of YF is characterized by several well-described phases or “periods” (*1*). During the “incubation period,” YFV replicates in the skin near the site of the inoculating mosquito bite, resulting in an initial burst of virus production and primary viremia that seeds YFV’s principal target tissue, the liver. YFV subsequently infects and kills hepatocytes resulting in a secondary viremia. During this “period of infection” individuals are acutely ill with non-specific signs (hyperemia, fever) and symptoms (malaise, headache, myalgia, nausea, epigastric pain, and vomiting). This is followed by a “period of remission” characterized by marked clinical improvement, and most individuals continue to recover without long-term sequelae. However, ∼30% of cases will relapse into a second sickening known as the “intoxication phase”, which is characterized by a return of fever, a distinct lack of viremia, multi-organ failure, neutrophilia, pancreatitis, and hemorrhagic manifestations such as black vomit (*1, 6, 12–14*). This phase of the disease is highly lethal and survivors are often left with significant sequelae. Despite centuries of observation, the development of “intoxication” in YF––its predisposing factors and pathophysiology––is poorly understood (*14*).

With few exceptions (*15*), the development of viscerotropic YF disease––*i.e*., a pattern of YFV infection that primarily involves the visceral organs––is limited to primate hosts (*16*). Rodents are naturally resistant to YFV infection and, when immunocompromised, exhibit a neurotropic pattern of infection without signs of viscerotropic infection or disease (*17*). Thus, the macaque monkey (*Macaca* spp.) has long been the gold standard animal model for studying YF pathogenesis (*1, 16*). However, the limitations inherent to high-containment nonhuman primate research have significantly hindered investigation of the mechanistic underpinning YF pathophysiology. Multiple YFV strains have been adapted to infect the Syrian golden hamsters (*Mesocricetus auratus*) via serial passaging, causing a viscerotropic pattern of disease characterized by hepatocyte destruction, elevations in the liver damage marker ALT, and the development of coagulopathy (*18, 19*). Although the hamster-adapted Asibi strain of YFV (HA-Asibi) has been used in preclinical evaluation of anti-YFV countermeasures (*20–24*), only a cursory examination of the disease caused by HA-Asibi in the hamster has been undertaken. Given the relative lack of small animal models of viscerotropic YF disease, we predicted that a detailed understanding of HA-Asibi pathogenesis in the hamster could be leveraged to advance our understanding of YF pathophysiology.

Here, we performed a detailed virological and pathological analysis of HA-Asibi in the Syrian golden hamster, which we then leveraged to examine early events in the development of severe YF. In particular, we observed that HA-Asibi infection causes severe intestinal ischemia despite a lack of YFV replication, resulting in breakdown of the gut mucosal barrier and translocation of luminal bacteria that tracks up the portal system, prompting a neutrophil-predominant leukocyte response. Re-examination of samples from severe human YF cases also revealed the presence of bacteria in the liver, with elevation of several plasma biomarkers indicative of intestinal damage and bacterial translocation. Thus, the intoxication phase of YF is a sepsis-like syndrome caused by translocation of bacteria from a damaged gastrointestinal tract, explaining several enigmatic aspects of intoxication-phase YF including black vomit, pancreatitis, and neutrophilia in the absence of detectable viremia (*14*).

## RESULTS

### Hamster-adapted (HA)-Asibi recapitulates key features of severe YF

We first performed a route × dose study to establish a dose of HA-Asibi that results in highly reproducible lethal disease. Intraperitoneal inoculation of 1×10^5^ focus-forming units (FFU) of HA-Asibi was the lowest dose that resulted in uniform disease kinetics, as determined by weight loss and a composite clinical score (**Table 1**), with significant disease developing between 5-7 days post inoculation (dpi) and a lethality rate of ∼80% by 8 dpi (**Figure 1A-C**). Strikingly, 100% of hamsters infected with HA-Asibi developed a petechial rash––a classic stigmata of intoxication-phase YF (**Figure 1D**). To understand the relationship between viral tropism and disease parameters in greater detail, we also performed a comprehensive virologic analysis in HA-Asibi-infected hamsters over time. Viremia consistently peaked at 3 dpi (**Figure 1E**). The liver had the highest concentration of viral RNA, followed by the spleen, organs of the cardiovascular and respiratory system, kidney, and the gastrointestinal tract (**Figure 1F**). Of note, viral loads in the brain were among the lowest observed for any tissue, demonstrating a lack of HA-Asibi neurotropism in the hamster, unlike what has been observed in mouse models of YFV infection (*17, 25*).

**Figure 1.**
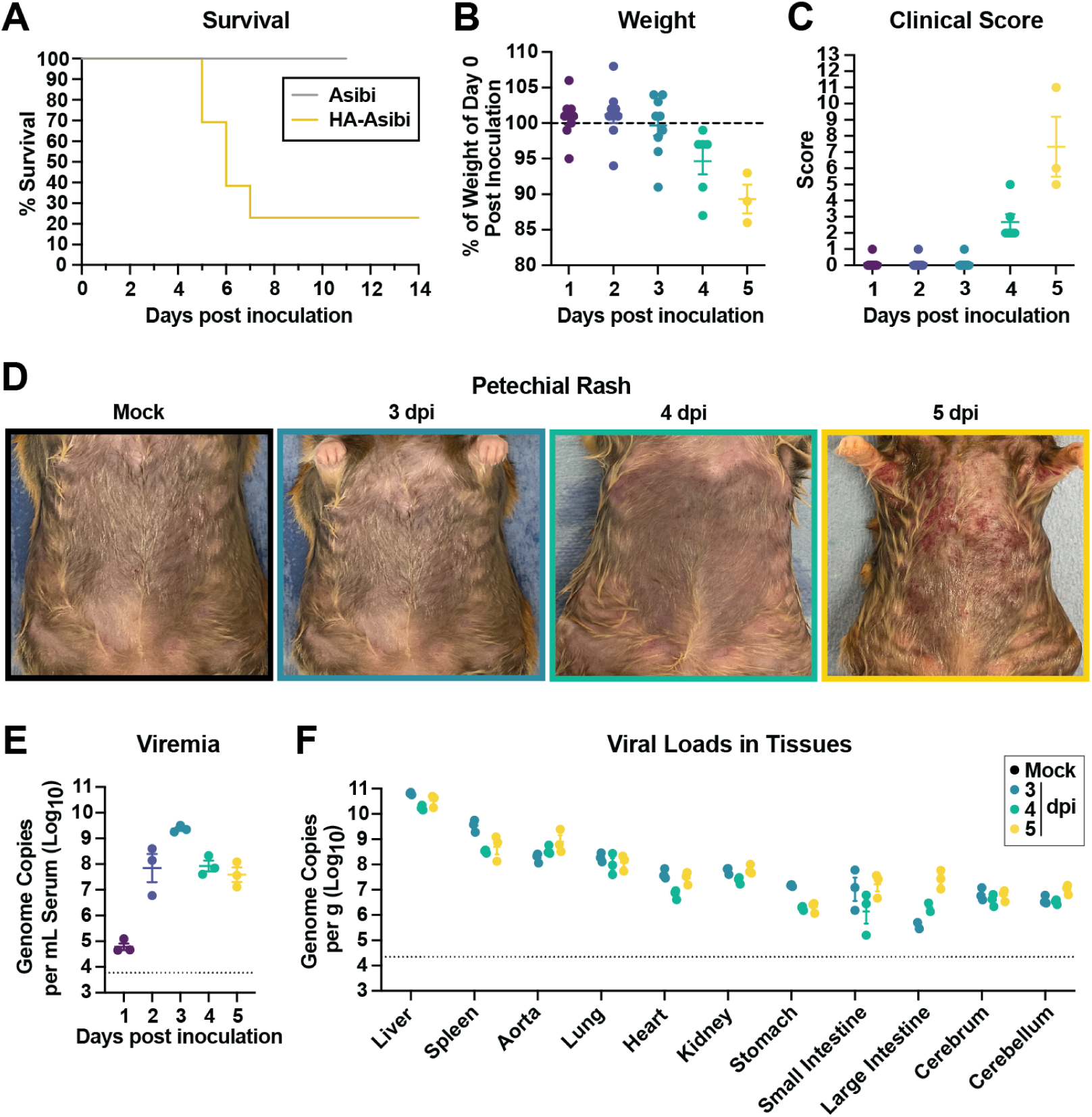
Hamster-adapted (HA)-YFV-Asibi causes lethal viscerotropic disease in hamsters. (A) Survival analysis of hamsters inoculated via intraperitoneal injection with 1×10^5^ FFU of wild-type (WT) Asibi (n=3) or hamster-adapted (HA)-Asibi (n=13); data is a composite of two independent experiments. (B) Percent weight change and (C) clinical scores following inoculation with HA-Asibi. Colors denote timepoints of 0/mock (black), 1 (purple), 2 (blue steel), 3 (teal), 4 (green), and 5 (yellow) days post inoculation (dpi) throughout the manuscript. (D) Development of petechial rash, shown here on the skin overlaying the chest, abdomen, and upper extremities. HA-Asibi viral loads in serum (E) and tissues (F) over the course of infection as determined by RT-qPCR. Dashed line shows the limit of detection; data are presented as mean ± standard error of mean (SEM).

### HA-Asibi causes severe hepatitis in hamsters

Serum concentrations of alanine aminotransferase (ALT, a biomarker specific to hepatocyte injury) became abnormally elevated starting at 3 dpi, with concentrations continuing to rise as infection progressed (**Figure 2A**). Evidence of gross liver injury was first noted on 3 dpi with the appearance of pale spots (mottling), which coalesced into larger patches of pale tissue by 4 dpi that affected the entire liver by 5 dpi (**Figure 2B**). Histologically, HA-Asibi induced many of the changes classically observed in human YF including destruction of the lobular architecture, microvesicular fatty changes (**Figure 2C,D**), and eosinophilic degeneration of hepatocytes (a.k.a., Councilman-Rocha Lima bodies) (**Figure 2E**). In late-stage disease a patchy multicellular infiltrate was observed in the livers of some HA-Asibi infected animals. We also observed patchy necrosis in the liver, as also seen in severe human YF. HA-Asibi-specific in-situ hybridization (ISH) showed widespread infection of hepatocytes throughout the liver (**Figure 2G**). Additionally, we expanded our ISH analysis to include several extrahepatic tissues, as infection of cells outside of the liver is presumed to occur but is poorly understood. However, with the exception of rare cells in lymphoid tissues, we did not identify any substantial population of extrahepatic HA-Asibi infected cells (**Figure S1**).

**Figure 2.**
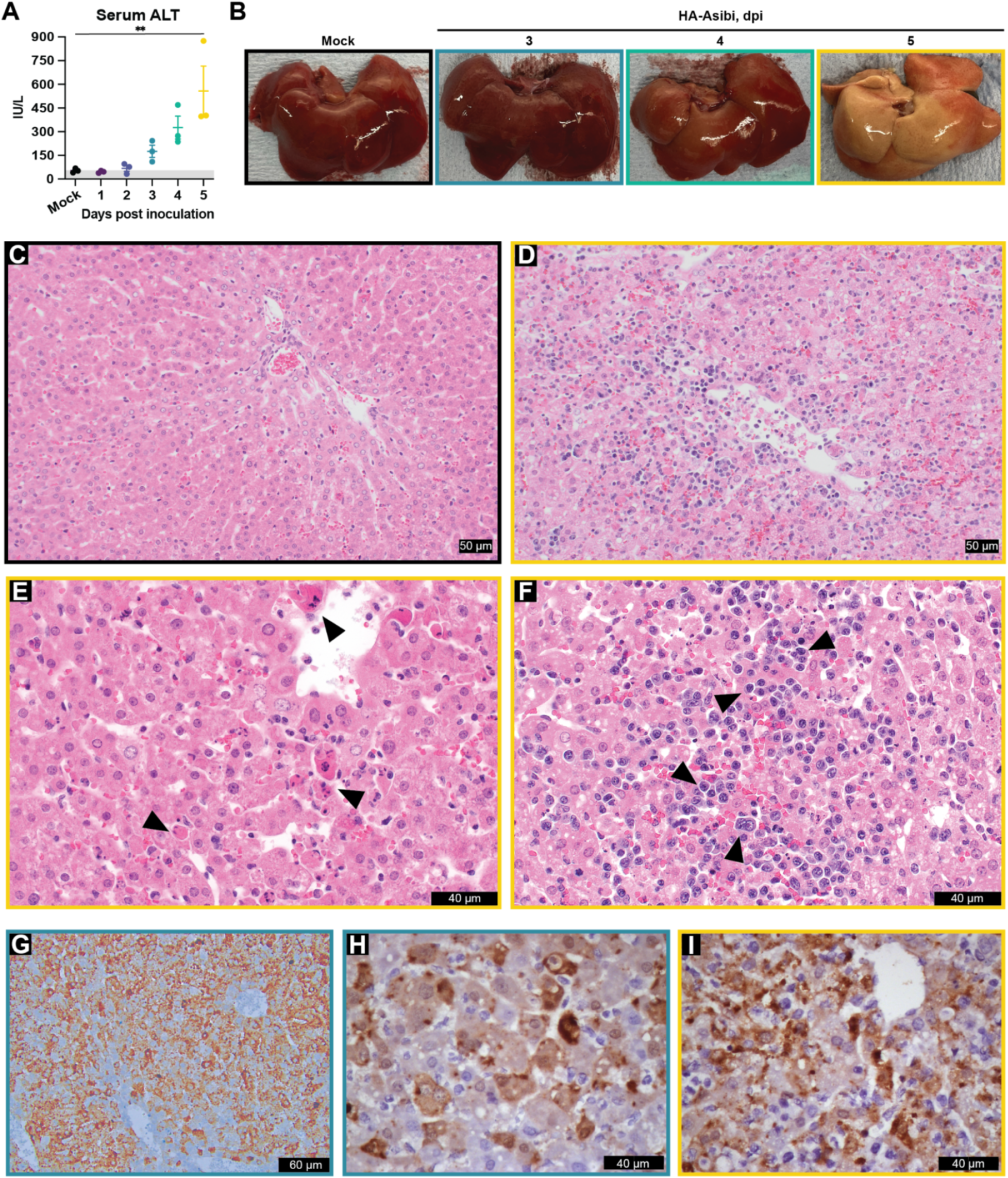
Liver pathology following HA-Asibi infection in hamsters. (A) Serum alanine aminotransferase (ALT) concentration, with human reference range in gray. Data are presented as mean ± SEM. Statistical significance determined from one-way ANOVA comparisons to mock-infected group (**: p<0.01). (B) Images of whole livers following HA-Asibi infection, illustrating gross pathology and pale/”shocked” appearance by 5 dpi. Hematoxylin and eosin (H&E) staining of FFPE liver tissue sections collected from (C) mock-(black outline) or (D-F) HA-Asibi-infected hamsters at 5 dpi (yellow outline). Upon infection, (E) Councilman-Rocha-Lima bodies (green arrowheads) representative of hepatocellular necrosis and (F) sinusoidal lymphoid/mononuclear infiltrates (black arrowheads) were observed. In-situ hybridization for YFV RNA (G) and immunohistochemistry for YFV antigen (H-I) in the liver showing pronounced hepatotropism.

### HA-Asibi infection results in global clotting factor insufficiency

To investigate whether infected hamsters develop coagulopathy similar to that observed in YF patients, we performed a series of clotting factor analyses. Prothrombin time (PT) and activated partial thromboplastin time (aPTT) are assays that evaluate the functionality of the extrinsic and intrinsic arms of the coagulation pathway, respectively **(Figure 3A)**. PT and aPTT began to prolong at 3 dpi **(Figure 3B**). While PT showed a slight prolongation at 4 dpi, aPTT greatly increased. By 5 dpi, both clotting times reached the upper limit of detection in most infected animals. To pinpoint the specific pathway(s) responsible for the defect in the coagulation system, we conducted mixing studies to evaluate the activity of clotting factors in the intrinsic (IX, VIII), extrinsic (VII), and common (X, II, I) pathways (**Figure 3C**). Over time, we observed a decline in the activity of all factors with the exception of fibrinogen––a well-known acute-phase reactant––which briefly increased from baseline levels before falling to abnormally low levels by 5 dpi.

**Figure 3.**
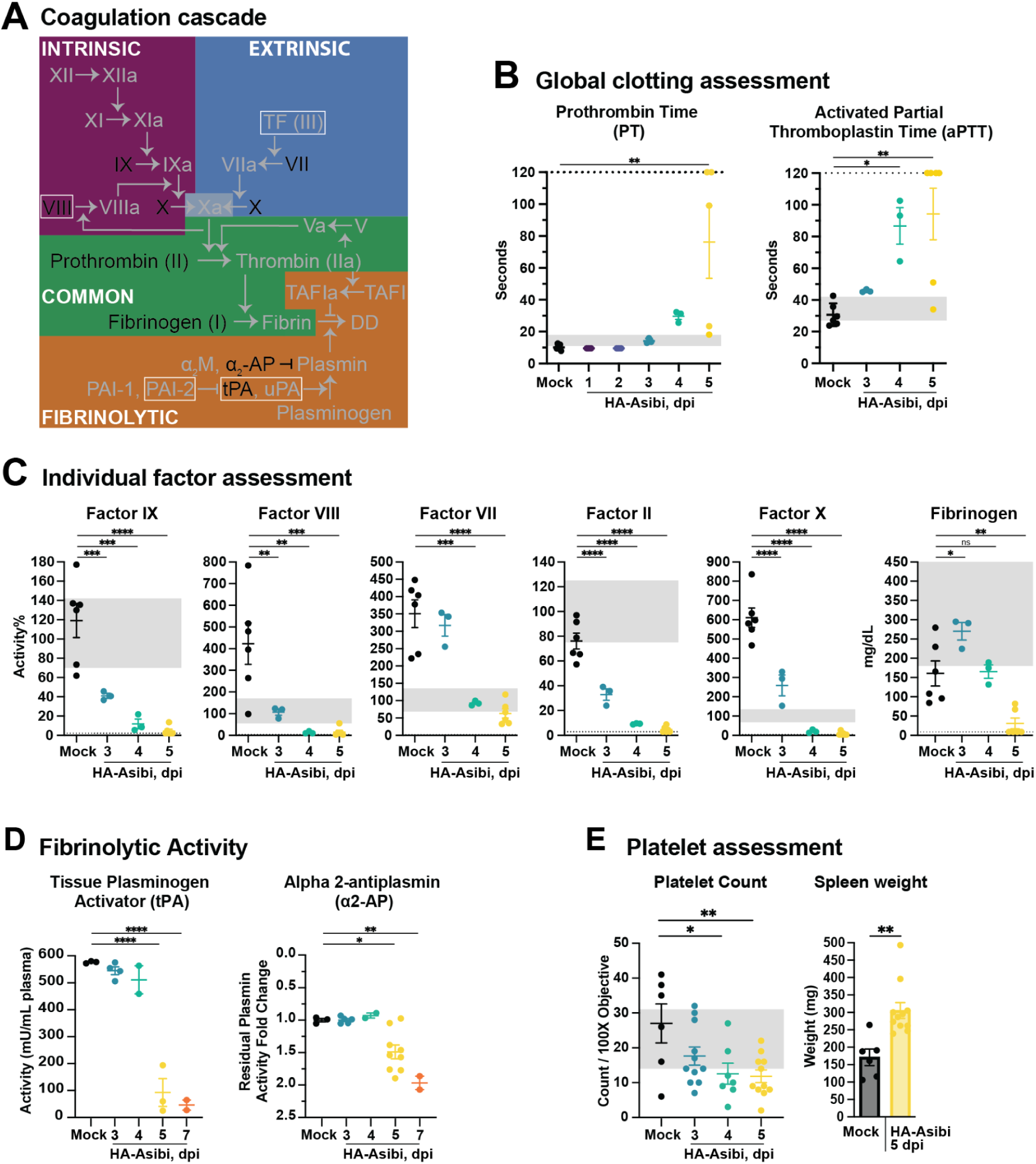
HA-Asibi infection induces coagulopathy defined by global depletion of clotting factors and coagulation regulators. (A) Simplified schematic of the coagulation cascade and the fibrinolysis pathway. Clotting factors and regulators assessed in this study are denoted in black, otherwise in gray. Boxes highlight factors not synthesized in the liver. Arrows and blunted arrows denote activation and inhibition, respectively. (B) Prothrombin time (PT) and activated partial thromboplastin time (aPTT) in hamsters upon HA-Asibi inoculation. (C) Coagulation factor activities determined using factor-deficient human plasma in PT-(VII, II, X, I) or aPTT-based (VIII, IX) mixing assays. (D) Tissue plasminogen activator (tPA) and ɑ2-antiplasmin (ɑ2-AP) activities determined with chromogenic substrate-based enzyme activity assays. For ɑ2-AP, data are normalized to the average of mock plasmin activity. ɑ2-AP activity is inversely proportional to residual plasmin activity. (E) Platelet count determined from blood smears and whole spleen weight of mock and HA-Asibi-infected hamsters at 5 dpi. Dotted lines represent the limits of detection of the assay; gray areas indicate the human reference range for each parameter as provided by the assay manufacturer. Data are presented as mean ± SEM. Statistical significance was determined from one-way ANOVA comparisons to the mock-infected condition and is denoted by an asterisk (*: p<0.05; = **: p<0.01; ***: p<0.001; ****: p<0.0001).

### Factor consumption plays a negligible role in coagulopathy induced by HA-Asibi

Clotting factor depletion can occur via two broad mechanisms: (1) defective clotting factor synthesis or (2) clotting factor consumption. Defective synthesis due to hepatocyte infection and destruction plays an obvious role in the development of coagulopathy in YF. However, fibrin split products (e.g., D-dimer)––indicative of a consumptive coagulopathy–– have also been reported in YF (*25, 26*). Unfortunately, a lack of assays for the detection of fibrin split products in hamsters is a known issue (*27*), and we were similarly unable to identify such an assay. Thus, we searched for other signs of clotting factor consumption including the presence of schistocytes (red cells that are sheared by intravascular fibrin strands) and the deposition of fibrin clots in the microvasculature of the kidneys, spleen, and lungs; however, we did not observe any of these features in hamsters infected with HA-Asibi (**Figure S2)**. We did identify a drop in platelet concentrations, and it appears that splenic sequestration may partially account for this observation, as evidenced by increased spleen mass over time (**Figure 3E**). During this body-wide survey, we identified surprising and extensive fibrin deposition in the pancreas, where catastrophic tissue damage led to the development of fibrinoid necrosis and saponification of the surrounding fat tissue by 5 dpi (**Figure 4**). Staining of the pancreas for YFV RNA by in situ hybridization confirmed a lack of YFV replication in this tissue (**Figure S1**), highlighting that tissue damage was not due to direct infection.

**Figure 4.**
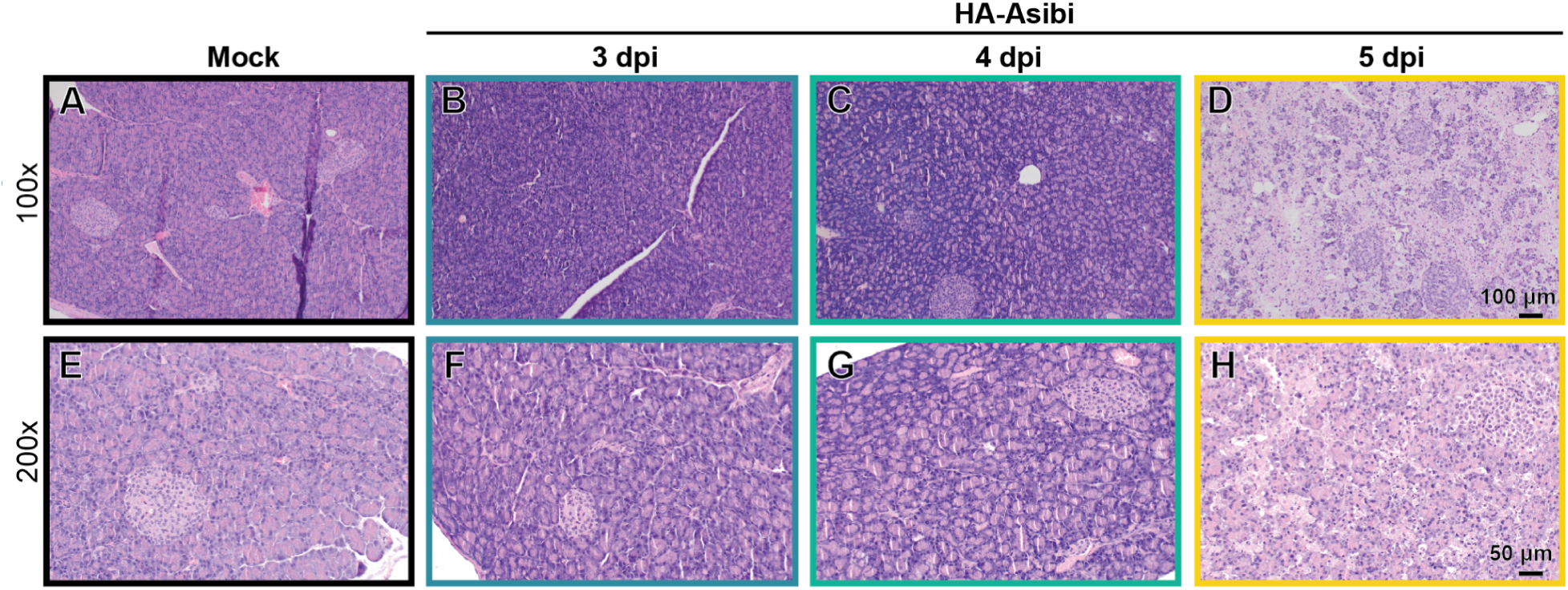
Pancreatic histopathology in HA-Asibi-infected hamsters. H&E staining of FFPE pancreas tissue sections collected from mock- or HA-Asibi-infected hamsters at 3-5 dpi. Images are representatives of each group.

### Intestinal damage precipitates bacterial translocation and sepsis in YF

The out-of-proportion tissue damage in the pancreas led us to further scrutinize histopathology in organs of the gastrointestinal system. Further investigation revealed that significant pathologic changes developed in the small intestine as HA-Asibi infection progressed. Gross changes included the development of edema by 3 dpi with the appearance of petechiae on the intestinal serosa by 4 dpi and black necrotic-appearing bowel by 5 dpi (**Figures 5A-H**). Histologically, this process began with edema and villous blunting at 3 dpi that progressed to focal hemorrhages in the villi and lamina propria by 4 dpi (**Figures 5I-L**). This was accompanied by separation of the villus epithelium from the lamina propria resulting in the formation of “Gruenhagen spaces”––a hallmark of ischemic injury (**Figures 5M,N**) (*28–31*). By 5 dpi, the epithelium at the tips of the villi had eroded, with coagulated blood and bacteria in and around the villi and crypts (**Figure 5O**). Given these histological changes, we wondered whether the mucosal barrier was functionally compromised and could serve as an entry point for bacteria and bacterial products. Indeed, by 5 dpi we were able to identify lipopolysaccharide (LPS)-positive material (indicative of Gram-negative bacteria) in the portal vein and liver parenchyma, along with LPS-laden histiocytes in the pancreas (**Figure 6A-C**). We also detected increased concentrations of serum endotoxin in the serum and liver of HA-Asibi-infected animals (**Figures 6D,E**) and were able to recover viable microorganisms from liver homogenates of HA-Asibi-infected animals (but not mock-infected animals) (**Figure 6F**). This was accompanied by the development of neutrophilia, which is typically seen in response to bacterial (but not viral) infections (**Figures 6G-I**). Collectively, these observations point toward a sepsis-like response in the intoxication phase of yellow fever, driven by intestinal barrier disruption and subsequent bacterial translocation.

**Figure 5.**
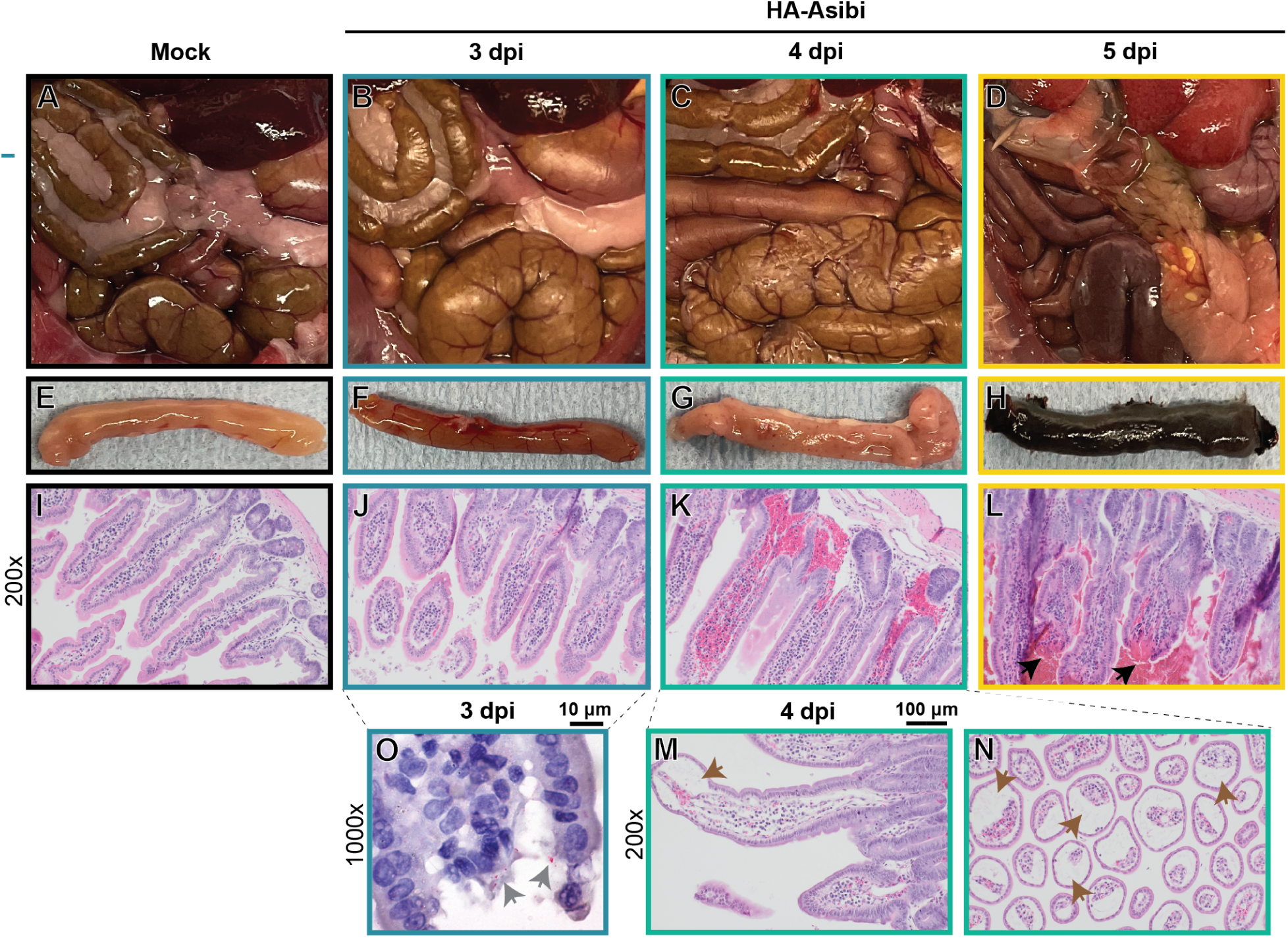
Gastrointestinal pathology in HA-Asibi-infected hamsters. (A-D) Images of gross small intestines illustrating development of (B) edema, (C) petechiae, (D) hemorrhage, and necrosis upon HA-Asibi infection. (E-H) Gross images of small intestinal segments illustrating development of hemorrhage and necrosis by 5 dpi at the time of euthanasia. (I-N) H&E staining of FFPE small intestine sections showing development of (I-L) hemorrhage and villous mucosal breakdown, and (M-N) “spaces of Gruenhagen” at tip of villi (brown arrows) indicative of ischemia. (O) Immunostaining of LPS (gray arrows) in the small intestine at 3 dpi. Images are representatives of each group.

**Figure 6.**
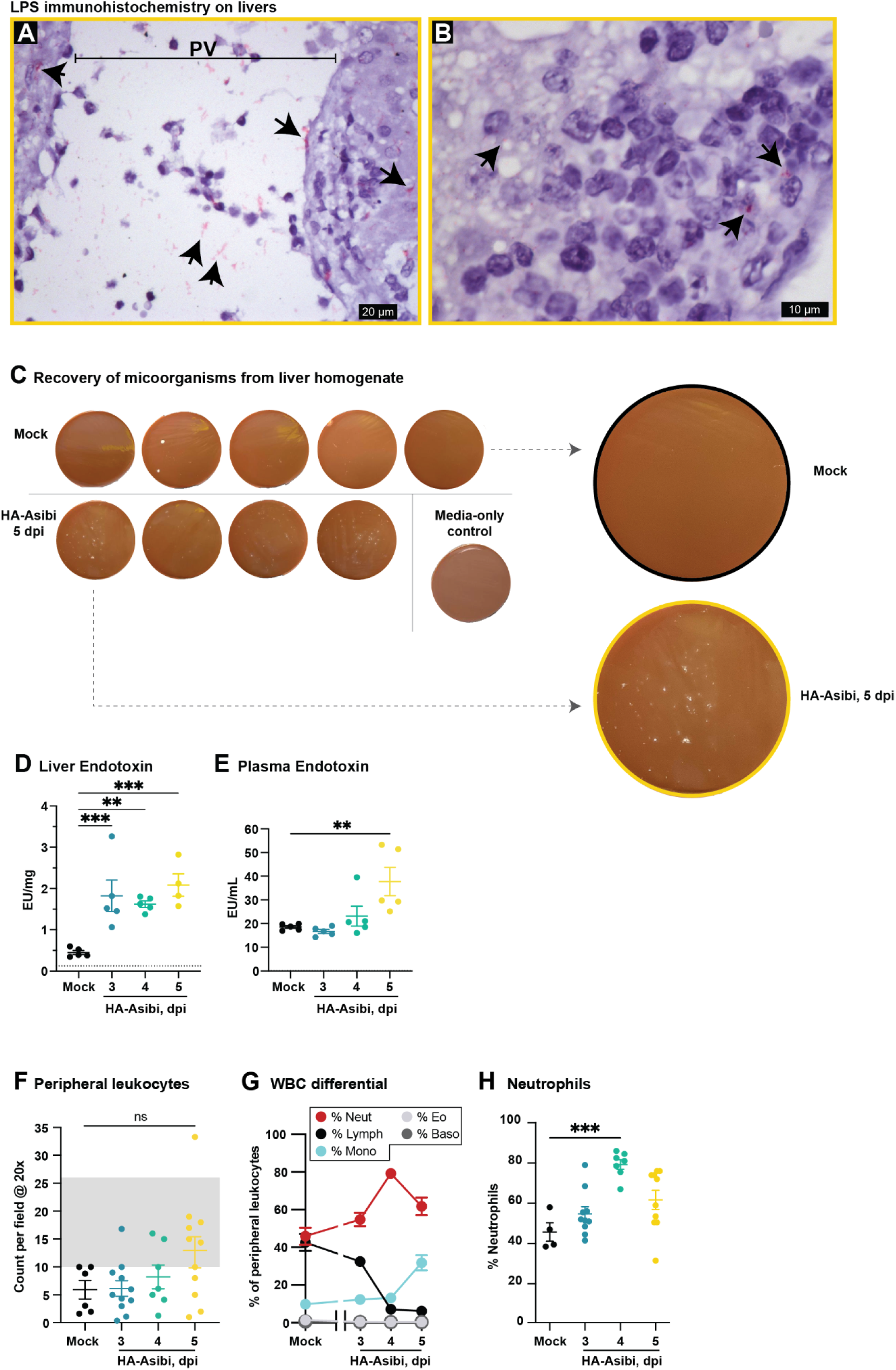
HA-Asibi infection leads to bacterial translocation. (A,B) Immunohistochemistry for bacterial lipopolysaccharide seen as pink material highlighted by arrows in portal vein (PV) and parenchyma of hamsters infected with HA-Asibi at 5 dpi. (C) Chocolate agar plates showing microbial growth from liver homogenates of mock/HA-Asibi-infected hamsters at 5 dpi. (D,E) Concentrations of bacterial endotoxin in the liver and plasma following HA-Asibi infection. (F-H) Total leukocytes, differential peripheral leukocyte count, and neutrophil count as determined by blinded assessment of peripheral blood smears after HA-Asibi infection. Data are presented as mean ± SEM. Dotted lines indicate the limits of detection of the assay; the gray area represents the human reference range as recommended by pathologists. Statistical significance was determined from one-way ANOVA and is denoted by an asterisk (*: p<0.05; = **: p<0.01; ***: p<0.001; ****: p<0.0001).

### Intestinal damage and bacterial translocation are correlates of YFV disease and lethality in humans

To determine whether a similar pathophysiologic process could explain YF intoxication in humans, we first examined gastrointestinal pathology in a collection of 17 human autopsy cases of severe YF from the 2017-2018 YF epidemic in the Metropolitan region of São Paulo, Brazil. Prior to death, all cases developed hallmark signs and symptoms of the intoxication phase of YF along with laboratory findings of transaminitis, neutrophilia, elevated serum levels of pancreatic enzymes (**Table S2**). Autopsies were performed following the Letulle technique with removal of all organs *en bloc* followed by dissection of each organ, formalin fixation, paraffin embedding, and H&E staining. Upon opening, all 17 cases showed gross abnormalities in gastrointestinal organs that included: dilated stomach with hemorrhagic wall; hemorrhagic necrosis of the pancreas; with steatonecrosis in the retroperitoneal fat tissue; dilated intestinal loops, and petechiae/hemorrhagic patches along the intestinal serosa (**Figure 7A,B**). Microscopic examination revealed vascular congestion, and endothelial damage (tumefaction, fibrinoid necrosis, and fibrin clots) along with interstitial edema and patchy hemorrhage throughout gastrointestinal tissues. In the small intestines, we observed ischemic changes and denudation of the epithelium, similar to what was observed in hamsters with late-stage disease (**Figure 7C, Figure S3**). We also found Gram-negative bacilli within small capillaries of the gastric mucosa and within the cytoplasm of macrophages in the gastric and intestinal mucosa (**Figure S4**). In addition to features of hepatitis characteristic of YF, we also found evidence for Gram-negative bacteria in the portal veins and parenchyma which was associated with a mixed inflammatory infiltrate (**Figure 7D, Figure S5**).

**Figure 7.**
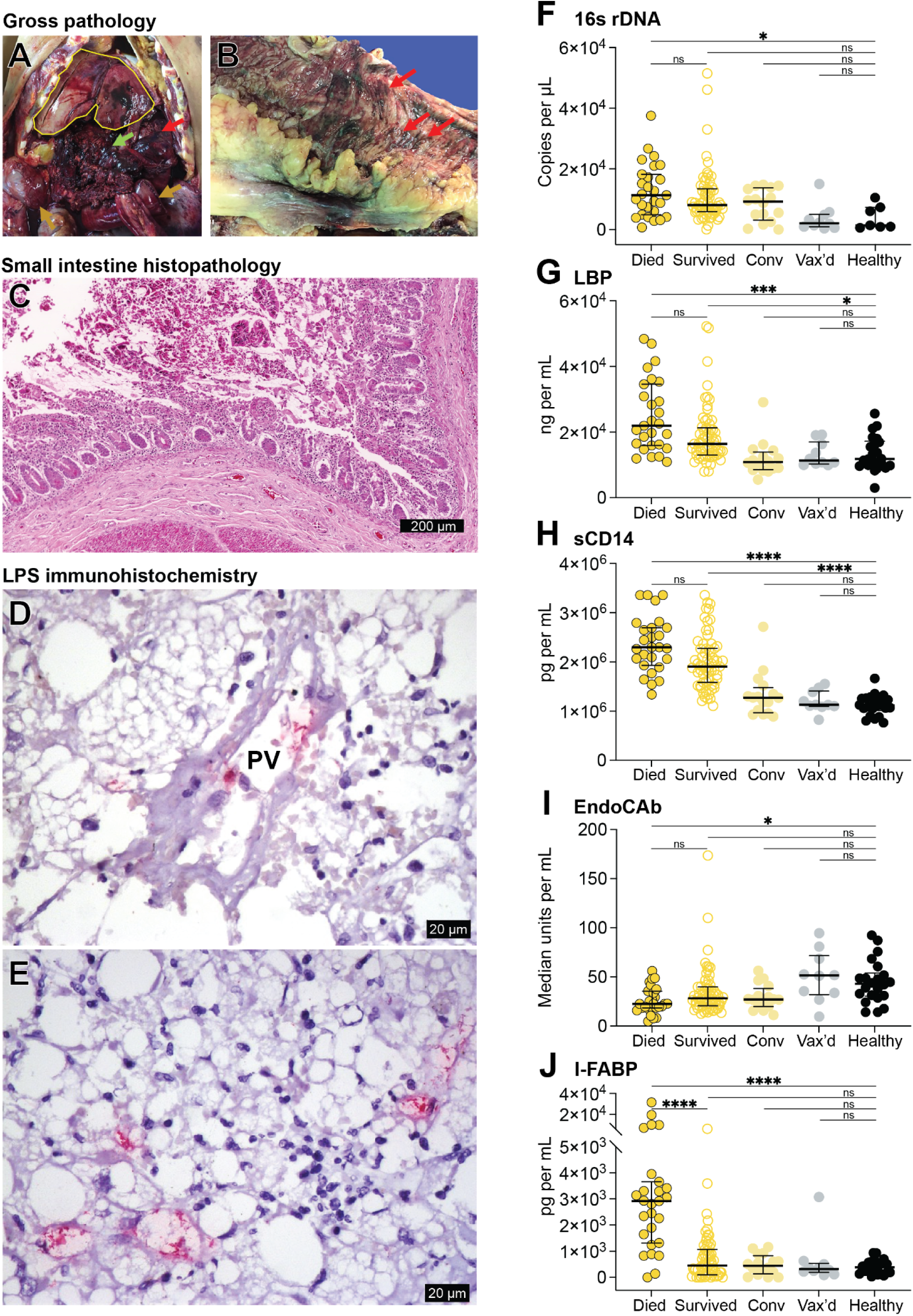
Intestinal damage and bacterial translocation are correlates of YFV disease and lethality in humans. (A) Abdominal organs after cavity opening in a 19-year old male YF victim showing enlarged liver with congestion and steatosis (yellow outline); dilated stomach with hemorrhagic wall (red arrow); hemorrhagic necrosis of the pancreas, with steatonecrosis in the retroperitoneal fat tissue (green arrow); dilated intestinal loops, with hemorrhagic walls (brown arrows). The abdominal cavity was filled with hemorrhagic fluid, mainly in the retroperitoneal area. (B) Ascending colon of a 47-year-old male YF victim showing confluent hemorrhagic spots (red arrow) and superficial ulcers on mucosa, associated with haemorrhages and steatonecrosis over the mesentery. The intestinal contents were hemorrhagic. (C) Hematoxylin and eosin stained small intestine from a YF victim showing epithelial denudation. (D,E) Detection of Gram-negative bacteria via LPS immunohistochemistry (IHC, red) in the portal vein and liver parenchyma of YF victims. Note the highly abnormal histological background due to YFV infection, showing degenerated hepatocytes and mixed tissue inflammatory reaction associated with Gram-negative bacteria. PV; portal vein. (F-J) Concentrations of 16s ribosomal DNA (F), Lipopolysaccharide binding protein (LBP) (G), soluble (s)CD14 (H), Endotoxin Core Antibody (EndoCAb) (I), and intestinal fatty acid binding protein (IFAB) (J), in samples collected at the hospital admission from patients who died from YF (n=27) or survived YF (n=63). As control groups, samples from survivors in the convalescent phase (Conv, n=16), YF vaccine (Vax’d, 14 days after vaccination) recipients (n=10) and healthy controls without recent vaccination (n=22) were used. Note, for the quantification of 16S rDNA, the number of healthy control samples tested (n=7) was lower due to the availability of samples. For all graphs, statistical significance was determined using the Kruskal-Wallis test with Dunn’s correction for multiple comparisons (*: p<0.05; **: p<0.01; ***: p<0.001; ****: p<0.0001). Only medically relevant comparisons are shown. Error bars show the median ± interquartile range.

We also undertook a quantitative biomarker analysis of our proposed mechanism of intoxication using a larger cohort of blood samples collected from the same 2017-2018 YF epidemic in Brazil, stratifying patients by outcome (death vs survival) and including a number of samples from relevant control groups (convalescent phase of YF survival, YFV-17DD vaccination, and healthy controls). First, we quantified bacterial 16S rDNA to directly assess the presence of bacteria in the blood, showing that fatal YF cases had significantly greater concentrations of plasma bacterial 16S rDNA compared to healthy controls (**Figure 7F**). We also examined several indirect markers of bacterial translocation including increases in LPS Binding Protein (LBP), decreases IgM endotoxin-core antibodies (EndoCAb-IgM), and increases in soluble CD14 (sCD14) as proxies for bacterial product translocation (*32, 33*), which also revealed statistically significant differences between fatal YF cases and healthy controls (**Figure 7G-I**). On average, these abnormalities were present but less significant in non-fatal YF cases, with a return to baseline in the convalescent group. Finally, we examined levels of intestinal fatty acid binding protein (I-FABP), a biomarker that is specific for intestinal damage (*34–36*). Plasma concentrations of I-FABP were significantly elevated in fatal YF cases relative to every other group including YF patients that survived, implicating gastrointestinal damage as a key differentiator between fatal and non-fatal YF (**Figure 7G-I**).

## DISCUSSION

In this study, we sought to understand the pathophysiology of the severe intoxication phase of YF using a combination of human autopsy data, human plasma samples, and prospective infection studies with a hamster-adapted YFV strain. Importantly, we found that the hamster model of YFV infection recapitulates several important features of the intoxication phase of YF disease in humans including hemorrhagic manifestations and high lethality. From these observations, we worked backwards in time to study early events in the development of the intoxication phase in the hamster model. Unsurprisingly, hepatitis was a major aspect of HA-Asibi disease. The development of coagulation abnormalities lagged behind liver damage, suggesting that the majority of the coagulopathy in YF results from a synthetic defect due to hepatocyte destruction. This was supported by coagulation factor mixing studies and the absence of histopathological features consistent with consumptive coagulopathy (*e.g*., fibrin deposition in tissues, red cell schistocytes). An exception to this was the significant fibrin deposition observed in the pancreas, likely secondary to ischemic necrosis. Pancreatitis is a recently recognized feature of severe YF in humans (*6, 13*); thus, it seems plausible that fibrinoid necrosis of the pancreas could explain the observations of elevated fibrin split products (*e.g*., D-dimer) in humans and nonhuman primates (*25, 26*).

In addition to the liver and pancreas, tissue damage was surprisingly severe across the gastrointestinal tract. An absence of viral RNA in these tissues indicated that this damage was not due to direct viral infection. Instead, the histologic changes in the small intestines––villous edema, endothelial damage, and the characteristic Gruenhagen spaces––,as well as those observed in gastric and colonic mucosa, were hallmarks of ischemic injury. Given that the pancreas and small intestine are both highly prone to ischemic injury, the severe damage to these organs is best explained by the development of ischemia in the splanchnic circulation, including mesenteric vascular bed (*28*). Notably, a correlation between I-FABP levels and intestinal damage resulting from ischemic processes is well demonstrated in humans (*35, 37*), and indeed we found significantly elevated levels of I-FABP in fatal human cases of YF, suggesting that measuring I-FABP may be a useful biomarker in caring for YF patients. Investigating the downstream consequences of YF-mediated intestinal damage showed that ischemia evolves into breakdown of the intestinal epithelium, which serves as a portal-of-entry for luminal bacteria that spread to the liver via the portal vein. This “second hit” likely impedes healing of the YFV-ravaged liver. Additionally, the reduced function of the liver as a filtering organ due to damage by YFV infection likely contributes to the spread of bacteria and bacterial products from the portal system into systemic circulation. Thus, the intoxication phase of YF appears to be a sepsis-like syndrome initiated by translocation of bacteria and bacterial products from the gastrointestinal tract.

While the upstream drivers of this ischemia may be multifactorial and remain to be determined, likely contributors may include hypovolemia due to vascular leakage, reduced fluid intake, losses via vomiting, reduced cardiac output (which is known to happen in YF (*i.e*., Faget’s sign and myocarditis (*38*)), and sympathetic/hypothalamic-driven splanchnic vessel constriction to maintain core blood pressure. The unique anatomy of the portal system, which collects venous blood from the organs of the gastrointestinal tract and delivers it to the liver, may also play a significant role in the development of mesenteric ischemia; acute hepatotoxicity (like that experienced in YFV infection) is known to elevate portal vein pressure which would provide additional resistance to the forward-flow of blood feeding the small intestines and pancreas (*39*). However, it is worth noting that severe gastrointestinal ischemia is not a feature found among other hepatitis viruses. Assessment of the factors that may contribute to gut ischemia in YF––in human patients or experimental animals––is difficult. However, a greater understanding of the pathophysiology of mesenteric ischemia in YF will likely yield further insights into how to treat and prevent this condition, such as with early and more aggressive fluid replacement.

Together, these findings help explain several perplexing features of YF disease. First, a unique feature of YFV infection relative to other viral hepatitides is the development of a transaminitis with concentrations of AST greater than ALT. Damage to the gastrointestinal tract is a potent source of AST, and it now seems likely that ischemic damage to the gastrointestinal tract is the source for high AST concentrations in YF. Second, the paradoxical findings of return of fever, lack of viremia, and presence of neutrophilia that define the intoxication phase of YF are entirely consistent with a secondary sepsis caused by the translocation of endogenous gut bacteria. Finally, this research brings renewed attention to the importance of the gut in YF pathogenesis. Although signs and symptoms of gastrointestinal disease such as nausea, hiccups, vomiting, abdominal distension, intense dorsal and epigastric pain, haematemesis (*i.e*., black vomit), and melena have historically been associated with severe YF (*40, 41*), more recent conceptualizations of YF pathophysiology have de-emphasized the role of the gastrointestinal tract, aside from the liver, in disease progression (*1*). We propose that the severity of mesenteric ischemia determines whether bacterial translocation––and ultimately relapse into intoxication––will develop in a YF victim. Understanding this disease process in greater detail will provide evidence for interventions to prevent and counteract the development of intoxication, with potential for reducing the mortality rate of YF.

## MATERIALS & METHODS

### Autopsy cohort

YF infection was confirmed in each case by at least one of the following methods: positive serum IgM; detection of YFV-RNA by RT-PCR in blood; or detectable YF antigen in tissues by immunohistochemistry. In all of the cases, infection with hepatitis A, B, C and D, dengue, zika, and chikungunya were excluded by serology and/or RT-PCR. Clinical and laboratory data from this cohort can be found in Table 1.

### Plasma samples and human subjects research

Ethical approval for the use of human samples in this study was obtained from the ethical review boards at the Instituto de Infectologia Emílio Ribas (CEP-IIER) and Hospital das Clínicas, University of São Paulo (CAPPesq) under ethical approval no. 6.741.440; CAAE n° 59542216.3.1001.0068; n° 0206/10; and ethical approval no. 3.190.194, CAAE n° 06704819.8.0000.0068. Consent for participation was obtained from all patients or from a legal representative when necessary, prior to inclusion in the study. All patient information was kept confidential throughout the study. The participants included in this study were recruited from two waves of a YFV outbreak in São Paulo state, Brazil, from January to May 2018, and from January to April 2019. A detailed description of the cohort characteristics can be found in two previous publications by our research group (*14, 42*). Briefly, we selected patients with confirmed YFV infection by one one the following methods: positive serum IgM; detection of YFV-RNA by RT-PCR in blood; or detectable YF antigen in tissues by immunohistochemistry. All individuals were older than 18 years, and with plasma samples available at the time of hospital admission.

### Animal use and Biosafety

All animal experiments were conducted in compliance with NIH and UW-Madison policies and procedures (IACUC: M006443; IBC: B00000929). Experiments involving wild-type and hamster-adapted yellow fever viruses were performed in ABSL-3 at University of Wisconsin-Madison by vaccinated personnel.

### Viruses for hamster studies

Wild-type YFV-Asibi was provided by the World Reference Center for Emerging Viruses and Arboviruses (University of Texas Medical Branch, Galveston, Texas). Asibi p7 strain was kindly provided by Dr. Alan D. Barrett (University of Texas Medical Branch, Galveston, Texas). A viral stock (designated as HA-Asibi in this study) was prepared from serum collected at 4 dpi after inoculation of a hamster with Asibi p7.

### Hamsters

Syrian golden hamsters (Charles River), female and 5-7 weeks of age, were randomized by veterinary staff upon arrival and housed in a controlled environment with regulated light/dark cycles, temperature, and *ad libitum* access to food and water in ABSL-3. Viral inoculations were performed via intraperitoneal (i.p.) injection of 100 µL containing 1×10^5^ focus-forming units (FFU) of virus diluted in tissue culture media (DMEM+2%FCS). Survival, weight, and clinical signs of disease were monitored daily. See Table S1 for clinical scoring rubric. Receiver operating characteristic (ROC) curve analysis of pilot survival studies showed that any animal with a clinical score of ≥8 would not survive, and this cutoff was used as criteria for euthanasia in subsequent studies. Blood was collected via gingival vein or via direct cardiac cannulation at the time of euthanasia. For euthanasia, animals were deeply sedated with ketamine via i.p. injection followed by thoracotomy.

### Quantitative YFV viral load by RT-PCR for hamster infection studies

At the time of euthanasia, animals were sedated and the cardiovascular tree was perfused with 50 mL of sterile saline. Tissues were then collected and snap-frozen. Tissue homogenization was performed by bead-beating in phosphate-buffered saline (PBS)+2%FCS using the Beadblaster-24 (Benchmark). RNA was extracted from tissue homogenates or serum using the MagMAX Viral Isolation Kit (Thermo Fisher, AMB1836-5) with the KingFisher Flex System (Thermo Fisher). RT-qPCR was performed using the TaqMan RNA-to-CT 1-Step Kit (Thermo Fisher, 4392653) on the QuantStudio 6 Pro Real-Time PCR System (Thermo Fisher) under the following cycling condition: 1) 48°C for 15 min, 2) 95°C for 10 min, 3) 95°C for 15 sec, 4) 60°C for 1 min, and 5) repeat from step 3 for additional 49 cycles. A primer-probe set for RT-qPCR (IDT) was designed that targets the 5’ UTR of YFV: forward primer (5’-AGGTGCATTGGTCTGCAAAT-3’), reverse primer (5’-TCTCTGCTAATCGCTCAAIG-3’), and probe (5’-/56-FAM/GTTGCTAGGCAATAAACACATTTGGA/3BHQ_1/-3’). An RNA standard was developed from a pCR™-Blunt-based construct (Invitrogen) that contains an 84-bp fragment from the 5’UTR region of YFV with the binding sites of the probe-primer set (5’-AGGTGCATTGGTCTGCAAATCGAGTTGCTAGGCAATAAACACATTTGGATTAATTTTAATCGTTCGTTGAGCGATTAGCAGAGA-3’). The construct was linearized with restriction enzyme digestion and *in vitro* transcription (MEGAscript T7 Transcription Kit, Invitrogen, AM1334) was performed. The transcript was purified (MEGAclear Transcript Clean-Up Kit, Invitrogen, AM1908), quantified, and diluted to 10^10^ copies/µL. Ten-fold dilutions of the transcript, ranging from 10^8^ to 10 copies/reaction, was used as a standard curve.

### Focus-Forming Assay for hamster infection studies

Vero-WHO cells were seeded at 5×10^4^ cells per well in a 96-well flat-bottom tissue-culture plate. Cells were cultured in DMEM (Gibco) supplemented with 5% fetal calf serum (FCS, Omega) at 37°C. The next day, medium was substituted with 100 µL of 10-fold serial dilutions of each virus-containing sample prepared in DMEM supplemented with 2% FCS. Cells were incubated at 37°C for 1 hour with rocking every 15 minutes. Subsequently, a 125-µL overlay of 2% methylcellulose (Sigma-Aldrich) in 2x MEM (Gibco) supplemented with 1% FCS was added to the cells. After 48 hours of incubation at 37°C, cells were fixed with 4% paraformaldehyde in PBS for approximately 25 minutes, washed with deionized water, and incubated in 100 µL of saponin-based permeabilization buffer containing anti-YFV 2D12 monoclonal antibody (1:800) overnight at 4°C with constant rocking. The antibody was prepared from clarified and filtered supernatant collected from mouse hybridoma cells (ATCC, CRL-1689). Cells were subsequently washed and incubated with goat anti-mouse secondary antibody conjugated with horseradish peroxidase (1:1000, Jackson ImmunoResearch, 115-035-062) at room temperature for 1 hour with constant rocking followed by TrueBlue peroxidase substrate (SeraCare, 5510-0030) for approximately 15 minutes. Foci were scanned and quantitated on a Biospot plate reader (Cellular Technology).

### Serum ALT assay for hamster infection studies

Serum ALT level was quantified using a commercial ALT reagent kit (Teco Diagnostics, A526-120) according to the manufacturer’s instructions with modification to a 96-well format. Briefly, 10 µL of serum was mixed with 50 µL of ALT substrate and incubated at 37°C for 30 minutes. 50 µL of ALT color reagent was added to the mixture. After a 10-minute incubation at 37°C, 200 µL of ALT color developer was added, followed by a 5-minute incubation at 37°C. The plate was read on a plate reader (CLARIOstar, BMG Labtech) at 505 nm.

### Coagulation factor assays for hamster infection studies

Prothrombin time was measured either from whole blood using the CoaguChek XS System (Roche) or citrated plasma using the Coatron X Pro (Teco) according to the manufacturers’ instructions. Activated partial thromboplastin time was measured using citrated plasma on the Coatron X Pro. Coagulation factor activities were determined using factor-deficient human plasma in PT-(VII, II, X, I) or aPTT-based (VIII, IX) assays.

### tPA Activity Assay for hamster infection studies

Tissue plasminogen activator (tPA) activity was determined using a colorimetric assay kit (Abcam, ab287878) according to the manufacturer’s instructions. In brief, citrated plasma from mock or HA-Asibi-infected hamsters was acidified with sodium acetate and diluted with an assay buffer. Acidified plasma samples were incubated with an inhibitor mix solution at 37°C for 10 minutes followed by addition of a substrate solution containing a serine protease-specific synthetic substrate that releases para-nitroaniline (pNA) chromophore upon cleavage. The inhibitor mix reduces the potential interference from other similar proteases, making the assay tPA-specific. Samples were immediately read for absorbance at 405 nm in kinetics mode (CLARIOstar, BMG Labtech) for an hour and tPA activity calculated based on a pNA standard curve.

### ɑ2-Antiplasmin Activity Assay for hamster infection studies

ɑ2-antiplasmin activity was determined using a colorimetric assay kit (Werfen, 0020009200). In brief, citrated plasma from mock or HA-Asibi-infected hamsters was incubated with human plasmin reagent at 37°C for 4 minutes followed by addition of a substrate solution containing a plasmin-specific synthetic substrate that releases pNA upon cleavage. Samples were immediately read for absorbance at 405 nm in kinetics mode (CLARIOstar, BMG Labtech) for 30min and residual plasmin activity calculated based on a pNA standard curve. ɑ2-antiplasmin activity is inversely proportional to residual plasmin activity.

### In situ hybridization for YFV in hamster infection studies

Viral RNA was visualized by in situ hybridization using a custom YFV-Asibi probe and the RNAscope 2.0 HD detection kit (Advanced Cell Diagnostics) according to the manufacturer’s instructions.

### Histopathology and peripheral smear evaluation

At the time of euthanasia, tissues were collected and placed in 10% neutral buffered formalin for fixation over 7 days. Hematoxylin- and eosin-stained formalin-fixed paraffin-embedded (FFPE) tissue sections and Wright-stained peripheral blood smears of mock and HA-Asibi-infected hamsters were examined by two trained pathologists Dr. Eduard Matkovic and Dr. Amaro Nunes Duarte-Neto. Histopathologic analysis of H&E tissues and evaluation of blood smears was performed using Olympus BX43 light microscope (Evident Corp.,Tokyo, Japan). Platelet count estimates from blood smears were obtained by counting the number of platelets in ten oil immersion fields using 100x oil objective, determining the average number per field, and multiplying the average number by 20,000 to produce an estimate per microliter. Validated white blood cells estimates were derived using a 40x objective and a multiplier of two. A manual 200-cell WBC differential count was performed on every slide.

### Immunohistochemistry (IHC)

A detailed IHC protocol for detection of YFV antigen was described previously (*43*). Briefly, after antigen retrieval in Tris-EDTA pH 9.0, the sections were incubated overnight with the primary antibody (goat-anti human IgM polyclonal, Institut Pasteur de Dakar, Senegal) at 1:20,000 dilution at 4°C. Slides were incubated with biotinylated secondary antibody (Reveal–Biotin-Free Polyvalent HRP DAB, Spring Bioscience, SPD-125) and chromogen (Dako Liquid DAB+ Substrate Cromogen System, Dako, K3460) and counterstained with Harris-hematoxylin (Merck, Darmstadt, Germany). The reactions followed standard protocols validated in our laboratories, using positive and negative controls. In all hepatic and intestinal samples IHC reactions were performed for detection of YFV antigens. The primary antibody was validated and tested in liver samples from patients with virus hepatitis (A, B, and C), herpes virus, cytomegalovirus, adenovirus, dengue virus, Treponema pallidum, Leptospira, and atypical mycobacteria infections, obtained from our autopsy archive. The IHC reaction was negative in all these samples. For the LPS IHC reactions, we performed antigen retrieval in citrate buffer at pH 6.0, with dilution of primary antibody (Ab35654 from Abcam) at 1:8000. A histological sample from a liver with cholangitis due to *E. coli*, which was confirmed by a bloodstream culture, was used as a positive control.

### Endotoxin Quantification in hamsters

The endotoxin levels in hamster plasma and liver homogenate samples were quantified using the Pierce™ Chromogenic Endotoxin Quant Kit (Thermo Scientific, A39552) following the manufacturer’s protocol. Briefly, plasma and liver homogenate samples were diluted 1:100 to match the kit’s detection limits and loaded into a 96-well microplate alongside endotoxin standards and blanks. Reconstituted amebocyte lysate reagent was added to initiate the reaction, and the plate was incubated at 37°C. After adding the chromogenic substrate, the reaction proceeded through a second incubation at 37°C, followed by termination with acetic acid. Absorbance at 405 nm was measured (CLARIOstar, BMG Labtech), and endotoxin concentrations were calculated based on a standard curve.

### Plasma biomarker assays for microbial translocation in human subjects

Quantification of microbial translocation-related plasma markers was performed using a commercially available Enzyme-linked Immunosorbent Assay (ELISA). The following markers were used: I-FABP (Human I-FABP, Hycult Biotech, HK406), to assess enterocyte death (intestinal tissue damage) and sCD14 (Human sCD14 Quantikine ELISA kit, R&D Systems, DC140), LBP (Human LBP, Hycult Biotech, HK315) and EndoCAb (EndoCab IgM, Hycult Biotech, HK504-IgM) as indirect markers of microbial translocation. All assays were performed according to the manufacturer’s instructions. The samples were diluted 200x for sCD14, 1000x for LBP, 10X for I-FABP and 50X for EndoCAb IgM. Absorbance readings were carried out using the Epoch Biotek spectrophotometer with Gen5 Data Analysis software. The concentration of each marker was determined by plotting a 4-PL logistic regression curve and adjusting the corresponding dilution factor to the final concentration of each marker.

### Quantification of Bacterial DNA from human plasma

Total DNA was obtained from plasma samples using the DNeasy Blood and Tissue kit (Qiagen, 69506) following the manufacturer’s recommendations, with an optimization step for better recovery of bacterial DNA, as previously described (*44*). Briefly, 500 μL of plasma was centrifuged at 5000 ×g for 10 min, and the material was split into three aliquots: 350 μL of the supernatant for one aliquot, and the remaining 150 μL was homogenized and divided into two aliquots of approximately 75 μL. DNA isolation from gram-positive bacteria was optimized by adding 180 μL of an enzymatic lysis buffer containing 20 mM Tris-Cl pH 8, EDTA (Sigma Aldrich) 2mM, Triton X100 1.2% and Lysozyme 20 mg/mL (Sigma Aldrich) to one of the 75 μL aliquots, followed by incubation at 37 °C for 30 min. The best recovery of gram-negative bacterial DNA was achieved by adding 180 μL of ATL buffer to an additional 75 μL aliquot. Following incubation, 20 μL of proteinase K and 200 μL of AL buffer were added into three aliquots of plasma, followed by incubation at 56 °C for 30 min (gram-positive bacterial aliquot) and 10 min for the remaining sample. The final steps were performed in accordance with the manufacturer’s instructions. Bacterial DNA was quantified using qPCR for the bacterial 16S ribosomal subunit gene, as previously described (*45*). To this end, we initially obtained a standard curve using *E. coli* DH5α (Thermo Fisher Scientific) pure culture as the source of genomic DNA. We spiked the genetic material in plasma samples from healthy donors to achieve similar conditions for the standard curve and biological samples to be tested. The final standard curve allowed quantification of 10^7^ to 10^2^ copies/μL, and the best qPCR reaction conditions were determined (Tables S3 and S4). Once the qPCR was standardized, quantification of the 16S DNA was performed on plasma samples from all YFV-infected patients in the acute (n = 90) and convalescent phases (n = 16), recently vaccinated individuals with 17DD (n = 10), and non-infected individuals (n = 7) as control groups. All samples were quantified in duplicate using the mean values for subsequent analyses.

### Microbial culture from hamster infection studies

Approximately ∼50 mg of tissue (liver or spleen) was collected with sterile instruments and transferred to tubes containing 400 µL of tissue culture medium (DMEM+2%FCS). Homogenization was performed for 20 sec using the Beadblaster-24 (Benchmark) with sterile beads, then 300 µL of homogenate was plated onto 85 mm chocolate agar plates (ThermoFisher) and incubated for 7 days, at which point plates were photographed.

### Statistical Analysis

Graphpad Prism 9 was used to perform statistical analyses. Details of each statistical test are reported in the figure legends.

## AUTHOR CONTRIBUTIONS

Conceptualization, ALB, XQ, ANDN, EGK; Methodology, MVT, CAC, MPM, ALB, XQ, ANDN, EM, ANDN, EGK; Investigation, all authors; Writing – Original Draft, XQ, ALB, MVT; Writing – Review & Editing, all authors; Funding Acquisition, ALB, MD, ANDN, EGK; Resources, ALB, ANDN, EM, MS, XQ, MD, EGK; Supervision, ALB, ANDN, EGK.

## SUPPLEMENTAL TABLES AND FIGURES

**Table S1.**
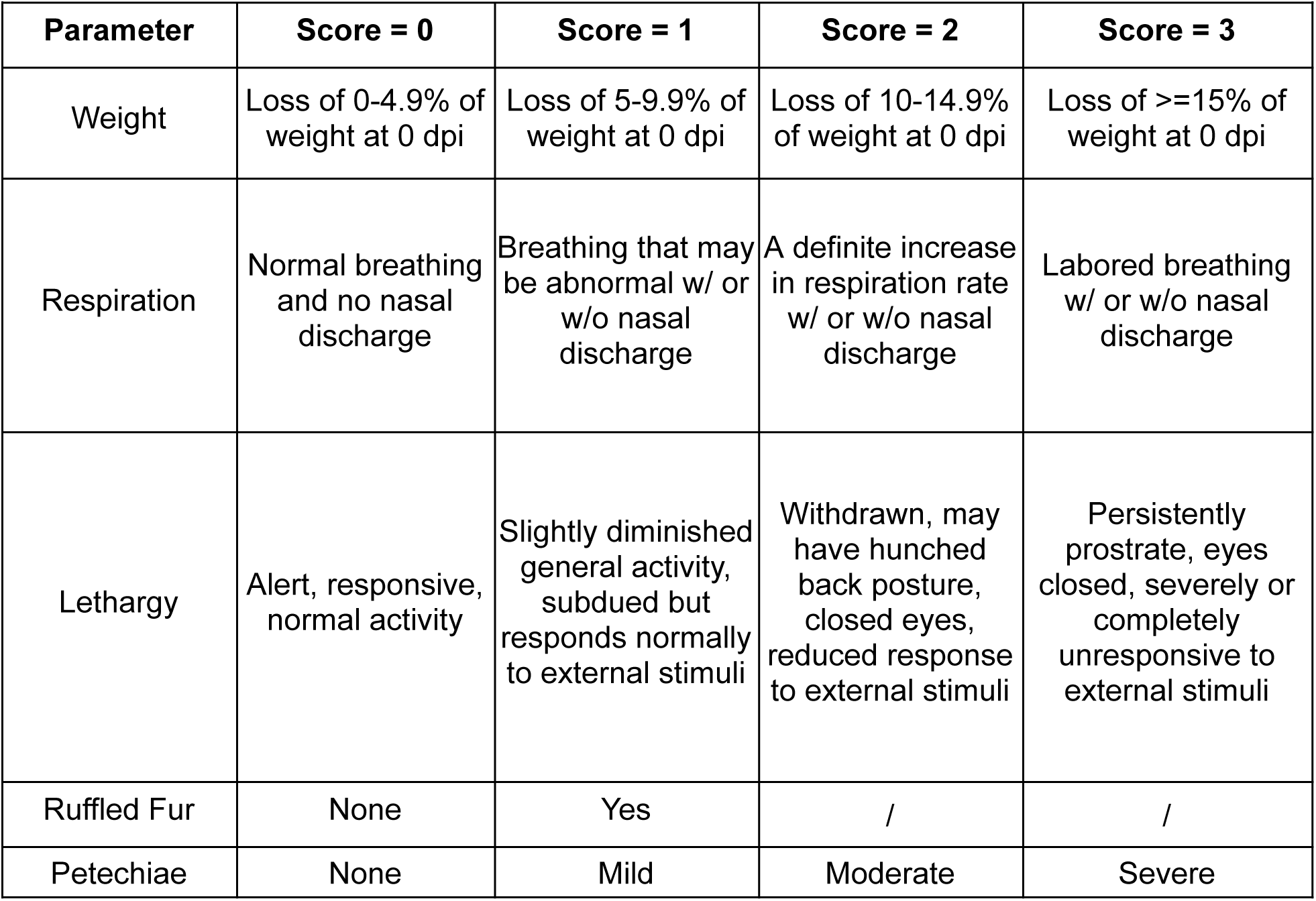
Clinical Scoring in Hamsters.

| Table S1. Clinical Scoring in Hamsters |  |  |  |  |
| --- | --- | --- | --- | --- |
| Parameter | Score = 0 | Score = 1 | Score = 2 | Score = 3 |
| Weight | Loss of 0-4.9% of weight at 0 dpi | Loss of 5-9.9% of weight at 0 dpi | Loss of 10-14.9% of weight at 0 dpi | Loss of $\geq 15\%$ of weight at 0 dpi |
| Respiration | Normal breathing and no nasal discharge | Breathing that may be abnormal w/ or w/o nasal discharge | A definite increase in respiration rate w/ or w/o nasal discharge | Labored breathing w/ or w/o nasal discharge |
| Lethargy | Alert, responsive, normal activity | Slightly diminished general activity, subdued but responds normally to external stimuli | Withdrawn, may have hunched back posture, closed eyes, reduced response to external stimuli | Persistently prostrate, eyes closed, severely or completely unresponsive to external stimuli |
| Ruffled Fur | None | Yes | / | / |
| Petechiae | None | Mild | Moderate | Severe |

**Table 2:**
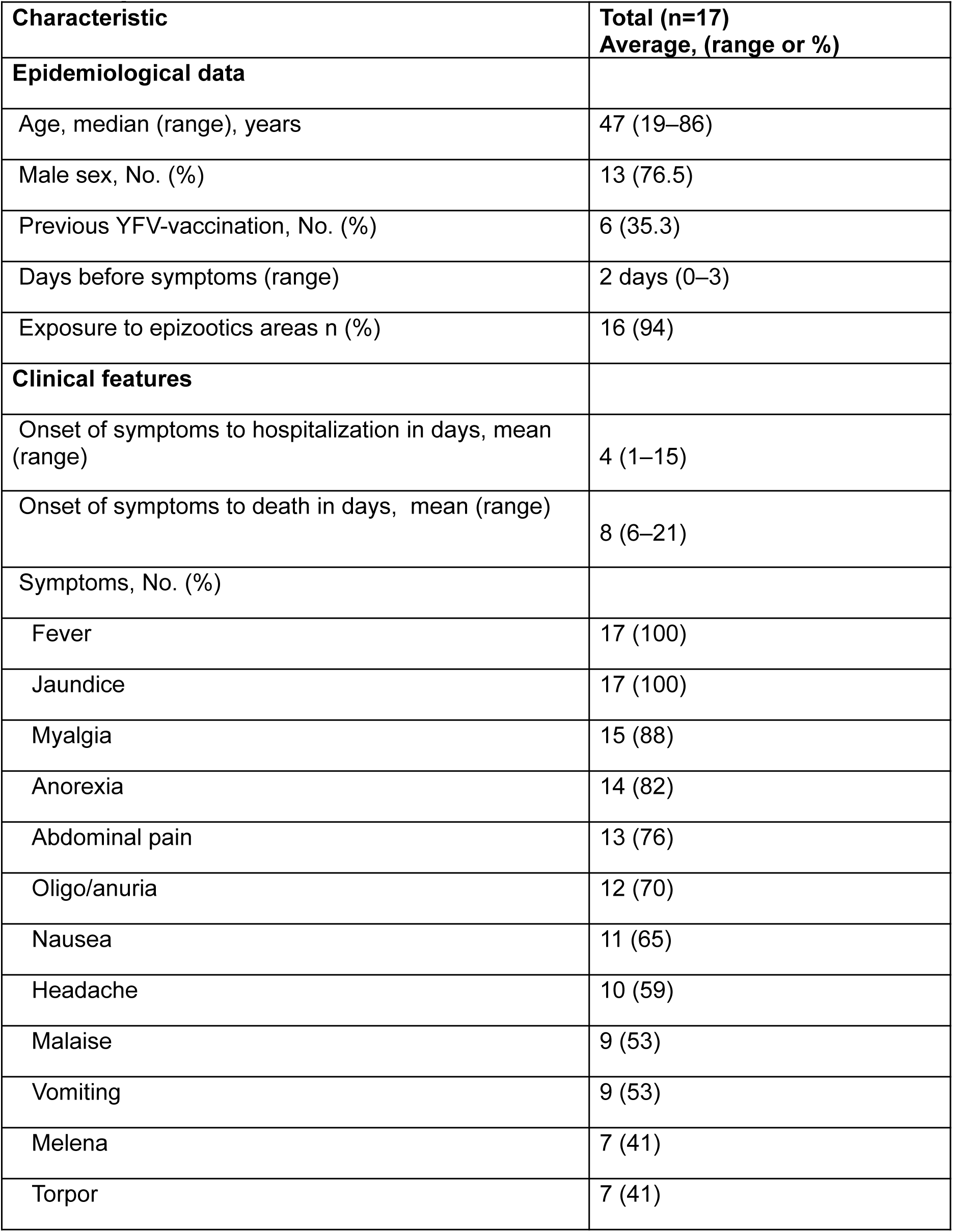
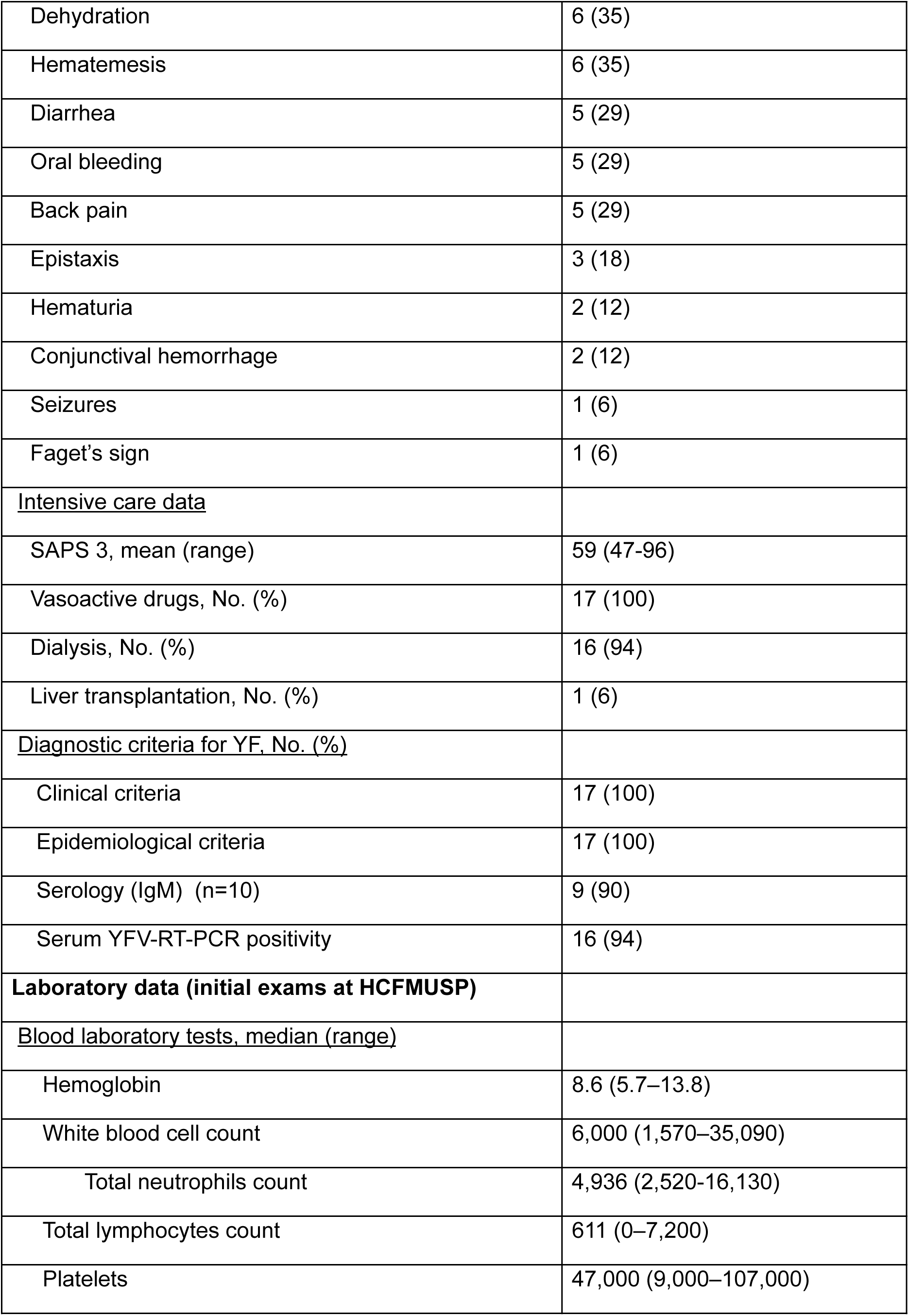
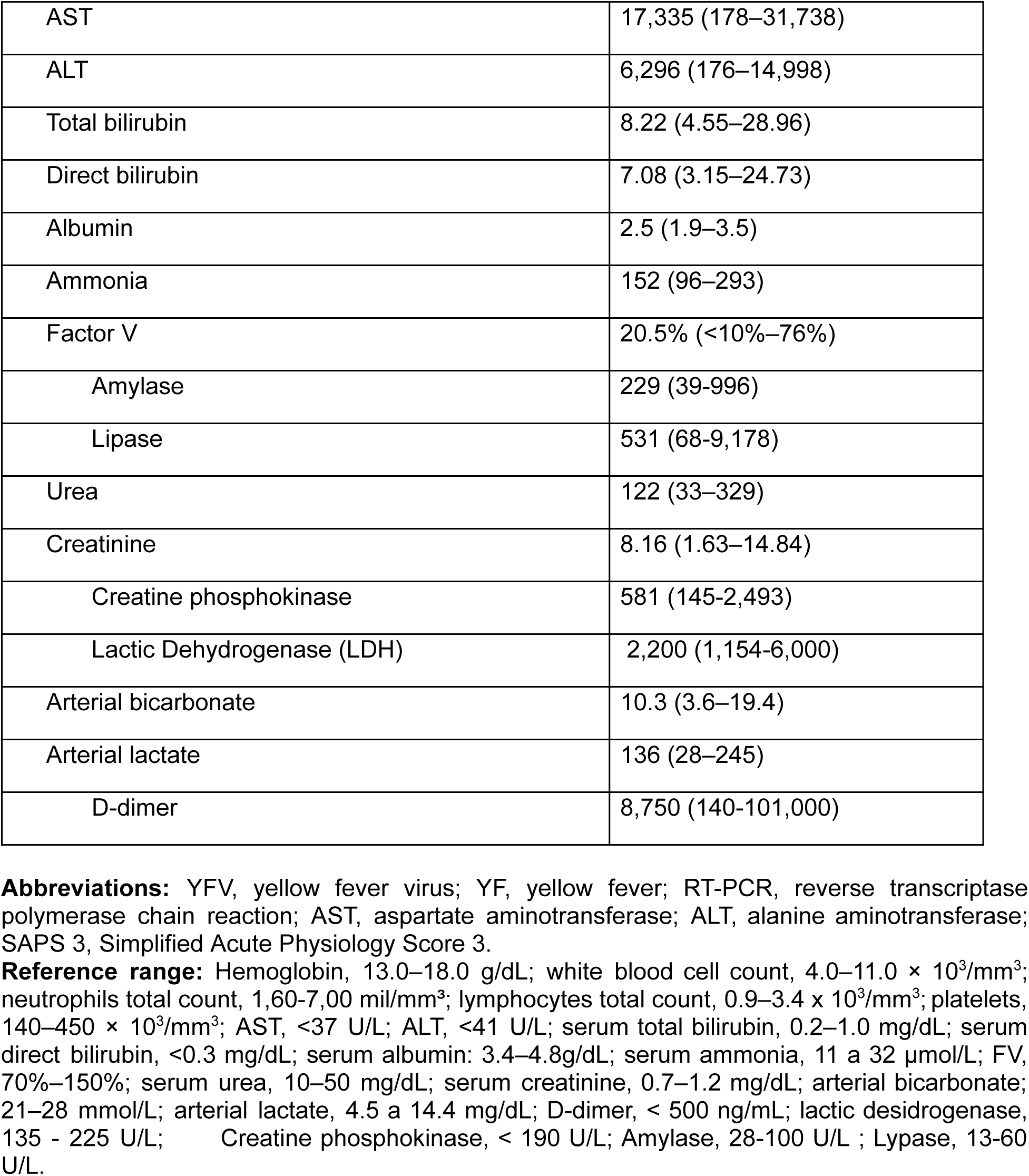
Epidemiological, clinical and laboratorial characteristics of 17 patients with confirmed yellow fever.

**Table S3.** PCR reaction for 16S DNA quantification.

| Reagent | Stock concentration | Use concentration | Volume per reaction (µL) |
| --- | --- | --- | --- |
| **TaqMan Fast Advanced Master Mix** | 2X | 1X | 10 |
| **Set02F** | 10 µM | 900 nM | 1,8 |
| **Set02R** | 10 µM | 900 nM | 1,8 |
| **Probe** | 25 µM | 200 nM | 0,2 |
| **DEPC water** | - | - | 4,2 |
| **DNA sample** | - | - | 2 |
| **Total** | - | - | 20 |

**Table S4.**
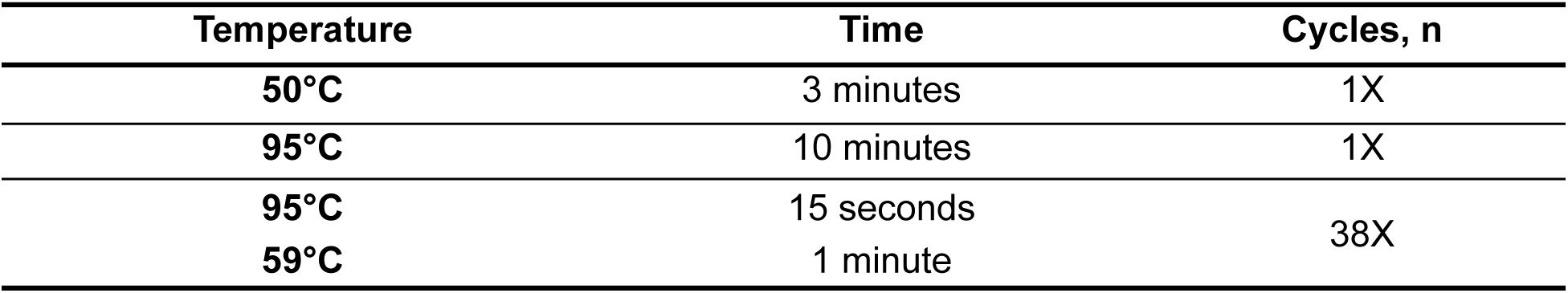
Thermocycling and amplification parameters.

| Temperature | Time | Cycles, n |
| --- | --- | --- |
| **50°C** | 3 minutes | 1X |
| **95°C** | 10 minutes | 1X |
| **95°C** | 15 seconds | 38X |
| **59°C** | 1 minute |  |

**Figure S1.**
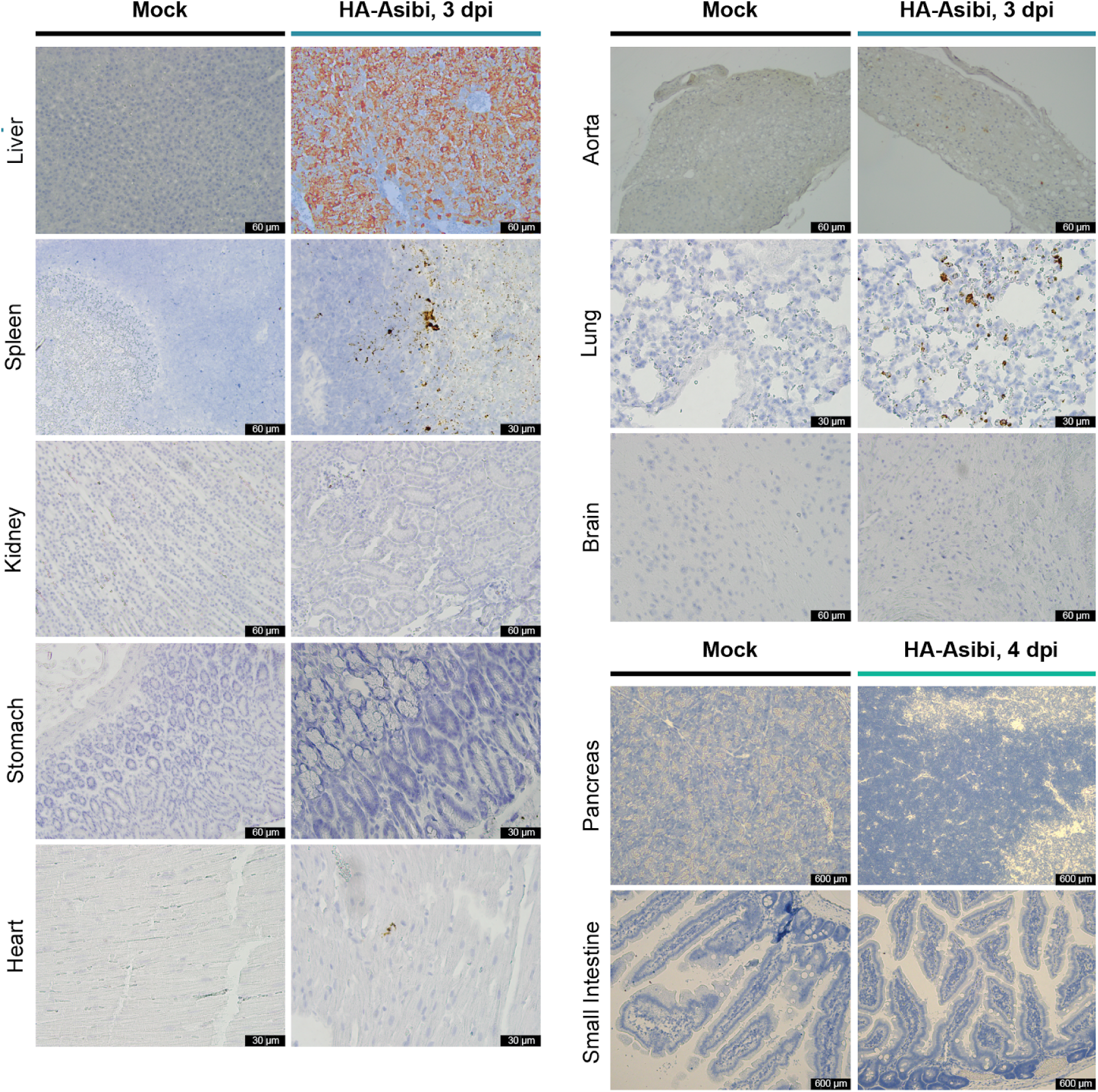
In-situ hybridization for YFV RNA in tissues from mock/HA-Asibi-infected hamsters.

**Figure S2.**
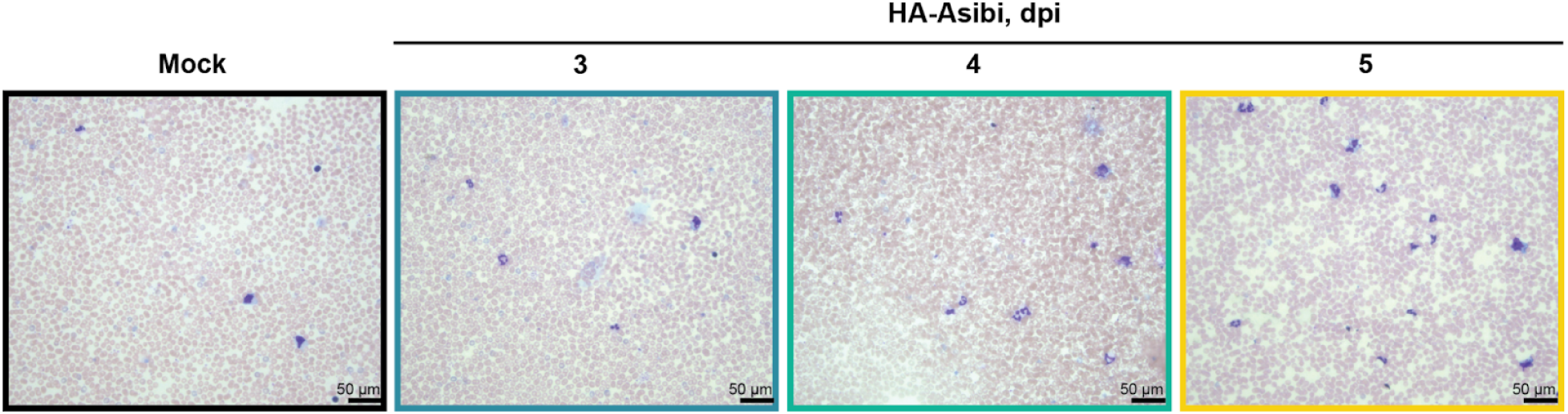
Peripheral blood smears from mock/HA-Asibi-infected hamsters.

**Figure S3.**
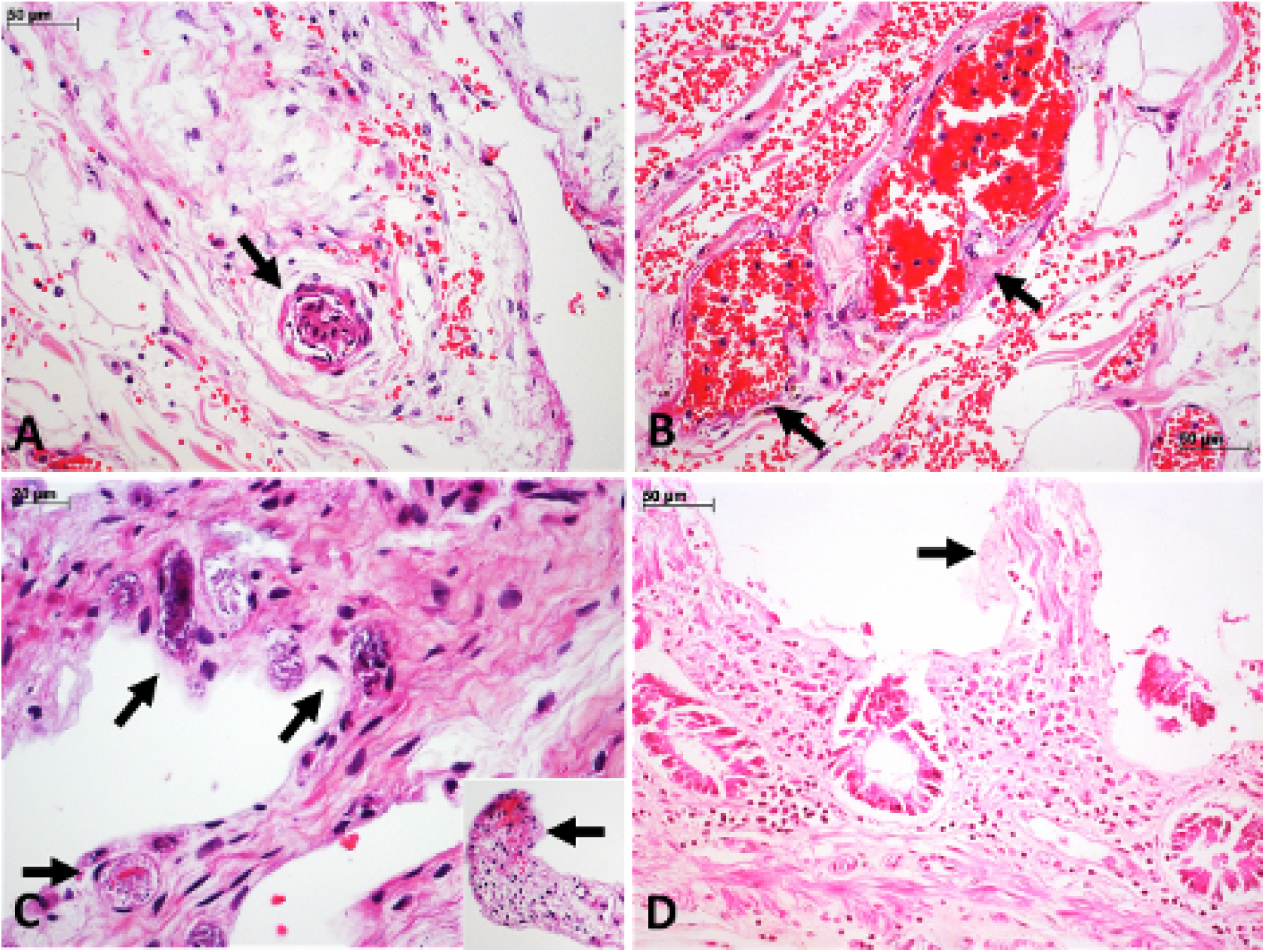
Pathological aspects of gastrointestinal tract among severe YF cases. A-B.Gastric submucosa with vascular congestion, interstitial edema and bleeding (A,B), with a small artery with a thrombus (A) and vessels with endothelial fibrinoid necrosis (B, arrows). C. Villi from small intestine exhibiting ischemia with epithelial denudation, and basophilic bacilli within small capillaries (arrows), and hemorrhage in the tip of an ischemic villus. D. Colonic mucosa exhibiting ischemic aspects, moderate inflammatory reaction in the lamina propria and epithelial autolysis.

**Figure S4.**
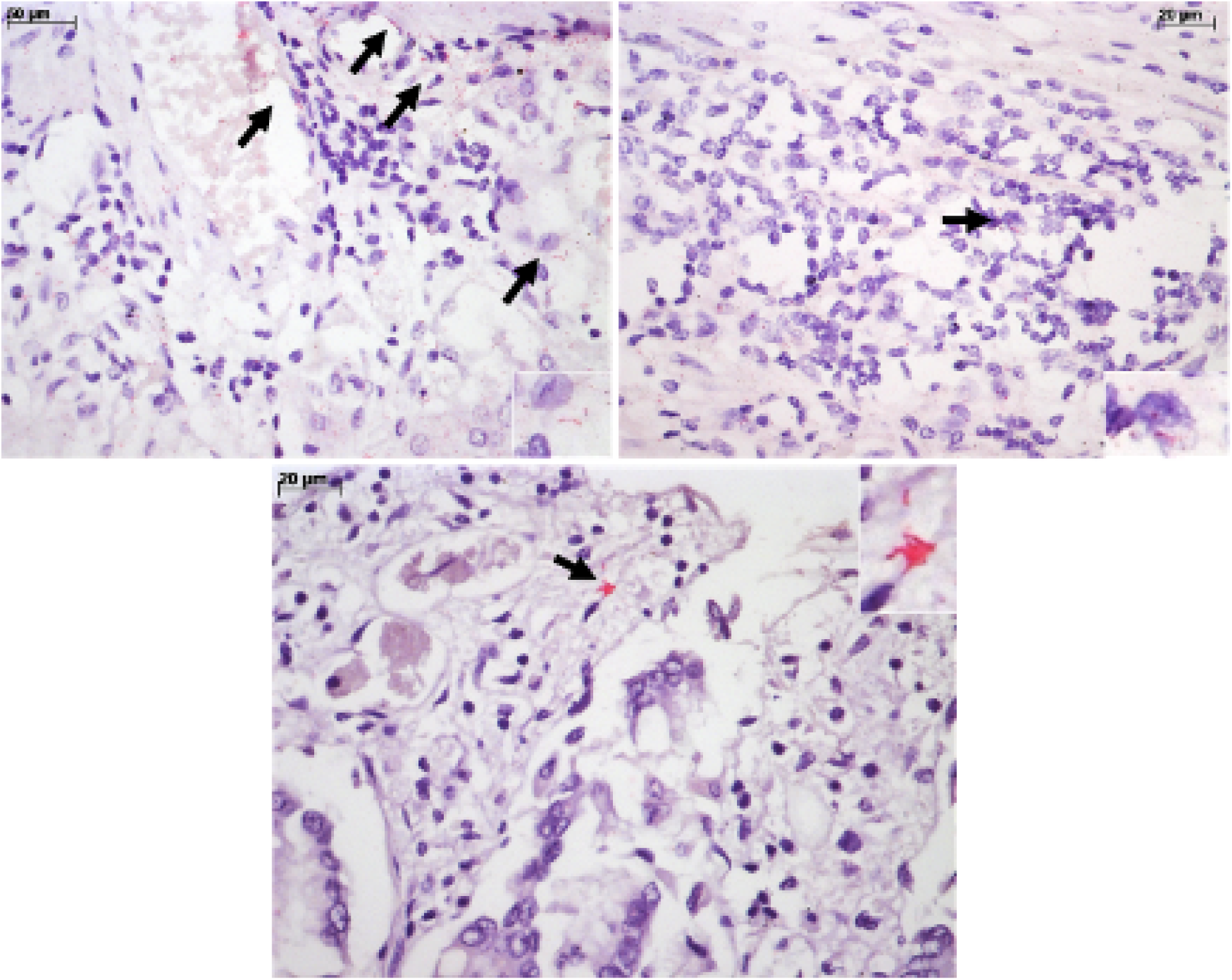
Gram-negative LPS detection in the gastrointestinal tract among severe YF cases. The LPS antigen is detected on the wall of Gram-negative bacteria, showing bacilli on within small capillaries of the gastric mucosa, within the cytoplasm of macrophages in the gastric and intestinal mucosa (A, B, insets) of bacilli in the ischemic colonic mucosa (C, inset). (LPS, IH, alkaline phosphatase).

**Figure S5.**
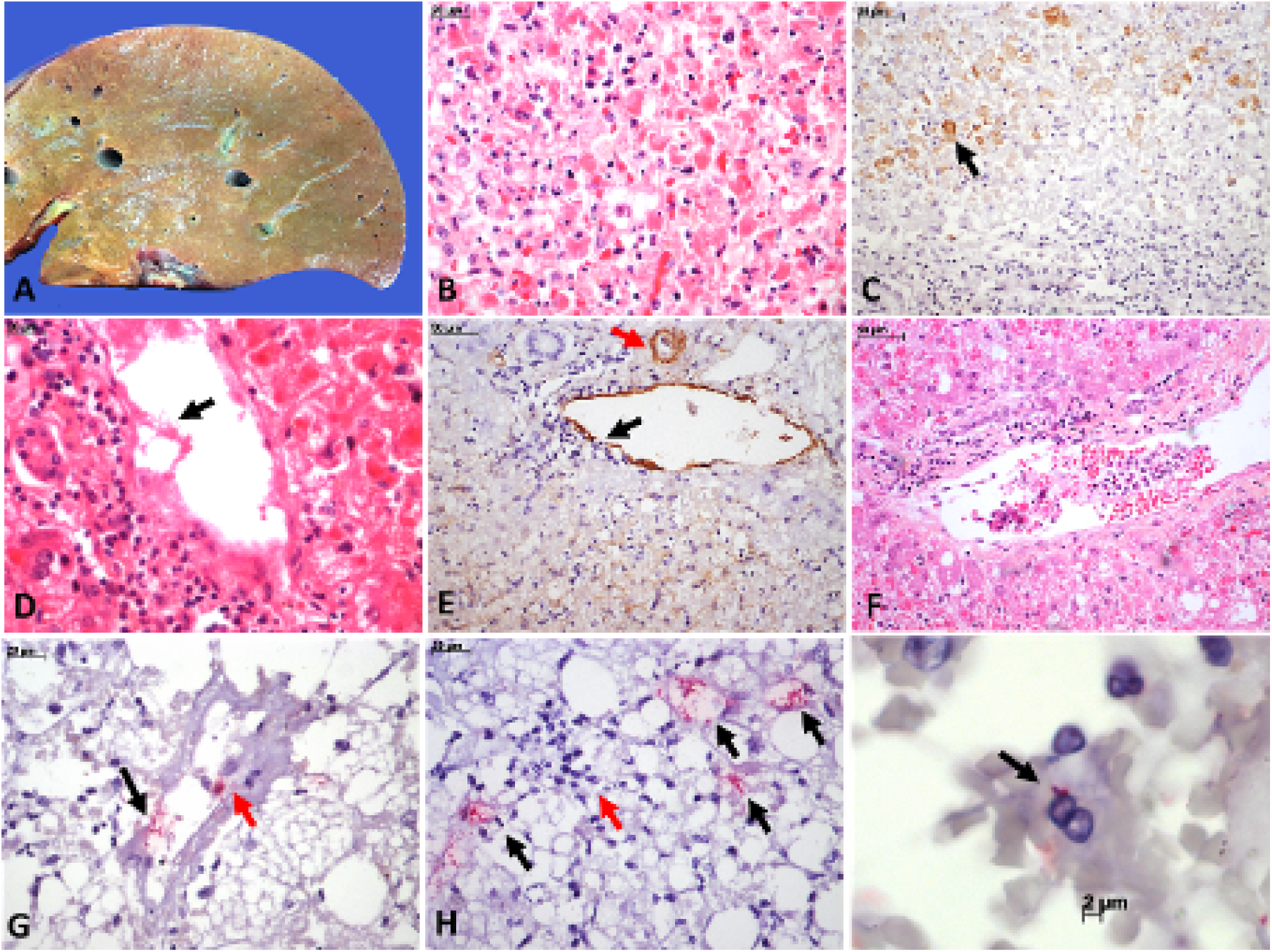
Pathological aspects of a human liver with YF fulminant hepatitis. A. YF-hepatitis macroscopy, showing diffuse steatosis and congestion. B-F. YF-hepatitis microscopic findings: steatosis and apoptosis of hepatocytes, hemophagocytosis by Kupffer cells, pyknotic bodies, mild inflammatory reaction by lymphocytes and some neutrophils in the mid-zonal area (B); YF-antigens expression in steatotic and apoptotic hepatocytes (C, immunohistochemistry, peroxidase); Portal vein exhibiting venulitis by mixed inflammatory reaction, and fibrinoid necrosis of endothelial line (D); Increased expression of VCAM in the hepatic artery branch and in the endothelial line of a portal vein exhibiting venulitis, and mild expression of VCAM in the sinusoidal endothelial cells (E); Portal vein with venulitis, leukostasis and fibrin aggregates (F); Presence of Gram-negative bacteria in a portal vein (G) and in the mid-zonal sinusoids, associated with inflammatory reaction (H), and attached to a sinusoidal lymphocyte (I) (LPS, IH, alkaline phosphatase).

